# RAB-35 and ARF-6 GTPases Mediate Engulfment and Clearance Following Linker Cell-Type Death

**DOI:** 10.1101/237289

**Authors:** Lena M. Kutscher, Wolfgang Keil, Shai Shaham

## Abstract

Clearance of dying cells is essential for development and homeostasis. Conserved genes mediate apoptotic cell removal, but whether these genes also control non-apoptotic cell removal is a major open question. <u>L</u>inker <u>c</u>ell-type <u>d</u>eath (LCD) is a prevalent non-apoptotic developmental cell death process with features conserved from *C. elegans* to vertebrates. Using microfluidics-based long-term in vivo imaging, we show that unlike apoptotic cells, the *C. elegans* linker cell, which dies by LCD, is competitively phagocytosed by two neighboring cells, resulting in cell splitting. Subsequent cell elimination does not require apoptotic engulfment genes. Rather, we find that RAB-35 GTPase is a key coordinator of competitive phagocytosis onset and linker cell degradation. RAB-35 binds CNT-1, an ARF-6 GTPase activating protein; removes ARF-6, a degradation inhibitor, from phagosome membranes; and recruits RAB-5 and RAB-7 GTPases for phagolysosome maturation. We propose that RAB-35 and ARF-6 drive an evolutionarily conserved program eliminating cells dying by LCD.

## INTRODUCTION

Clearance of cells undergoing programmed cell death is important during development of multicellular organisms, and failure to remove dying cells is implicated in developmental abnormalities^1^ and autoimmune disease^2^. Studies of the nematode *C. elegans* uncovered genes required for engulfment and degradation of cells dying by apoptosis^3–8^. Homologous genes regulate apoptotic cell clearance in *Drosophila* and vertebrates, as do a number of species-specific genes^9–11^.

While caspase-dependent apoptosis is a common cell death mode, recent studies demonstrate that caspase-independent non-apoptotic cell death processes are equally relevant in development^12^ and in disease^13^. While the molecular and genetic characterization of some non-apoptotic cell death processes has advanced considerably, whether common clearance mechanisms are used for apoptotic and non-apoptotic dying cells remains a major open question.

LCD (<u>l</u>inker <u>c</u>ell-type <u>d</u>eath) is a non-apoptotic developmental cell-death process that is morphologically conserved from *C. elegans* to mammals^12^. This cell death process is characterized by nuclear envelope crenellation, lack of chromatin condensation, and swelling of cytoplasmic organelles. These hallmarks of LCD are prevalent in vertebrate developmental settings, including degeneration of the Wolffian and Müllerian ducts during female and male gonadal development, respectively^14–16^, and motor neuron elimination during spinal cord formation^17^. LCD morphology is also characteristic of dying cells in several disease states, including striatal neuron death in Huntington’s disease^18^.

During *C. elegans* development, the male-specific linker cell leads the elongation of the developing gonad, and eventually dies by LCD^19^. Linker cell death is independent of caspases and all known apoptosis, necrosis, and autophagy genes. A network of proteins governing linker cell death has been uncovered, converging on the stress-induced transcription factor HSF-1, acting in a stress-independent manner to promote expression of LET-70/UBE2D2, an E2 ubiquitin ligase. LET-70/UBE2D2, together with other components of the ubiquitin proteasome system, then drive LCD and linker cell clearance^20–22^. Following LCD initiation, the linker cell is engulfed^23^. The linker cell, therefore, is a particularly attractive in vivo model for investigating the clearance of a cell that normally dies non-apoptotically: the time of linker cell death onset is predictable, the process can be followed in live animals, and engulfment events can be dissected genetically.

Here we show that engulfment and degradation of the linker cell differs in mechanics and genetics from apoptotic cell clearance. We demonstrate that, unlike apoptotic cell corpses, the linker cell is simultaneously engulfed by two U cell-descendent phagocytes, resulting in cell splitting. Apoptotic engulfment genes are not required for this novel form of engulfment. Rather, we find that the GTPase RAB-35, not previously implicated in apoptotic corpse removal, is a key coordinator of at least two steps in linker cell degradation. Early on, RAB-35 localizes to extending phagocyte pseudopods, and prevents premature onset of phagocytosis. Once engulfment has occurred, RAB-35 drives degradation of the linker cell by promoting recruitment of RAB-5 and then RAB-7 GTPases onto phagosome membranes, and subsequent lysosomal fusion and degradation. We demonstrate that both activities of RAB-35 require inactivation of another conserved small GTPase, ARF-6, which we show functions as a clearance inhibitor. RAB-35 physically interacts with the ARF-6 GTPase activating protein CNT-1, providing a plausible mechanism for ARF-6 inhibition. Furthermore, while RAB-35 localizes to engulfing-cell and phagosome membranes during most of linker cell clearance, ARF-6 only transiently persists at the membrane, and its timely removal depends on RAB-35.

Our work establishes a powerful in vivo model to study phagocytosis of cells that die non-apoptotically in development, and suggests that different phagocytic pathways can target cells that die by different means.

## RESULTS

### The Dying Linker Cell is Simultaneously Engulfed by Two Neighboring Cells

Linker cell death and clearance occurs during the transition from the fourth larval stage to the adult, and can take up to 8 hours to complete^19,24^. To examine this process at high spatiotemporal resolution, we used a long-term imaging microfluidic device we previously developed^24^ to simultaneously image the linker cell (*mig-24*p::Venus) and the engulfing cells (*lin-48*p::mKate2) in live animals (Figure 1A-B). Animals were loaded into the device, and z-stacks were acquired every 8 min for > 20h (Figure 1A). Animal development and linker cell death kinetics were not affected by either imaging or fluorescent reporter identity (Figure 1B, Table S1, top, see also^24^. The timings of characteristic events accompanying linker cell degradation were noted, with time zero corresponding to the time of first contact between the linker cell and the U.lp or U.rp engulfing cells (Figure 1C, Movie S1), and are consistent with previous reports^19,24^.

**Figure 1.**
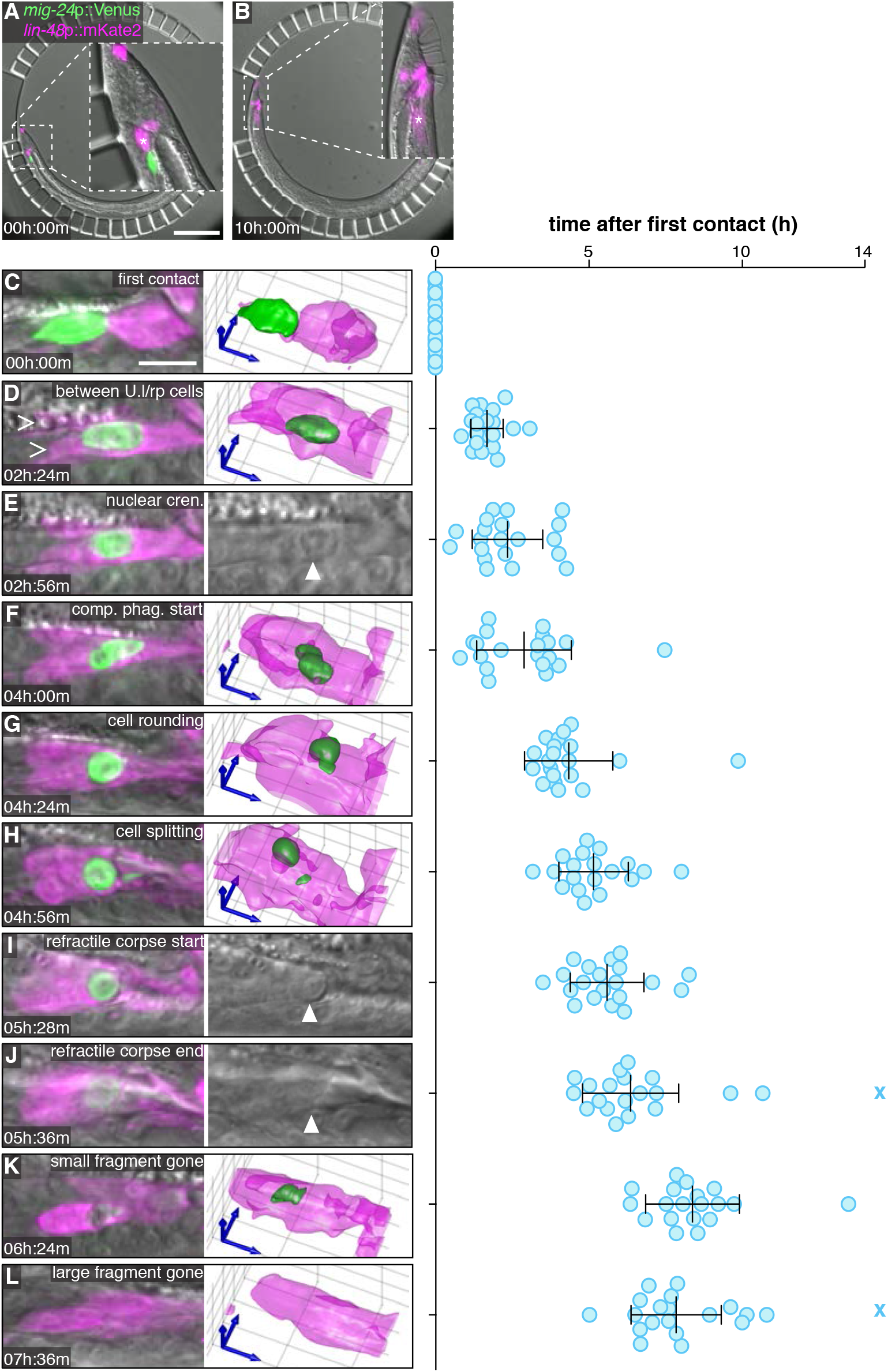
Linker Cell Death and Degradation. (A,B) *C. elegans* male immobilized in a microfluidic chamber (see 24) at (A) first contact between linker cell (green) and engulfing U.l/rp cell (magenta, asterisk), and (B) after 10 hours. All males examined in all figures carry the male-producing *him-5*(*e1490*) or *him-8*(*e1489*) mutation. Scale bar, 100 μm. (C) Left: Maximum intensity projection showing linker cell at first contact with U.l/rp cells. Right: 3D rendering. Scale bar, 10 μm. (D-L) Stereotypical events during linker cell dismantling. Images as in (C), except that in (E,I,J) a single DIC slice is shown instead of 3D rendering. Right, time of events in individual animals (blue circles) after first contact. X, event did not occur during imaging. Bars, mean ± sd. Arrowhead, linker cell. (D) Linker cell between U.l/rp cells (caret) (E) Nuclear crenellation onset. (F) Competitive phagocytosis begins. (G) Linker cell rounding (H) Linker cell corpse splits. (I) Onset of refractility. (J) End of refractility. (K) Small fragment disappears. (L) Large fragment disappears.

We found that after first contact, the linker cell migrates so that it becomes sandwiched between the U.lp and U.rp cells (Figure 1D), whose cell nuclei are displaced anterior to the linker cell. Linker cell nuclear crenellation also becomes apparent at this time (Figure 1E). Remarkably, following linker cell nuclear changes, both U.l/rp cells simultaneously attempt to engulf the linker cell (Figures 1F; S1A-D). During this process, which we term *competitive phagocytosis*, linker cell rounding (Figure 1G) is followed by splitting of the linker cell into two fragments (Figures 1H; S1E,F). The larger linker cell remnant contains the nucleus, and is equally likely to be found within U.lp or U.rp (8/19 and 11/19, respectively; p=0.5, χ^2^-test). The cell engulfing the nucleus-containing fragment is more likely to be binucleate (17/19; p=0.0006, χ^2^-test), a result of an earlier unrelated U cell descendent fusion event^23^, suggesting that the larger U.l/rp cell engulfs a larger portion of the linker cell. The larger linker cell remnant, but not the smaller, then becomes refractile by DIC microscopy (Figure 1I,J). The two linker cell fragments are degraded with similar kinetics (Figure 1K,L). Competitive phagocytosis does not result in extensive leakage of linker cell cytoplasm, as we never observed reporter protein outside the cell (N=35).

To identify the precise time point at which the linker cell fragments become internalized, we followed linker cell death in animals expressing the mKate2-PH reporter, derived from PLC-δ1, which marks the plasma membrane and membranes of open phagosomes by binding PI(4,5)P_2_^25^. Localization of this reporter around the linker cell ceases soon after cell splitting (Figure S1G-J), suggesting that linker cell fragmentation and U.l/rp internalization are coupled.

The unique mechanics governing linker cell dismantling raise the question of whether U cell descendants are specifically equipped for linker cell clearance. To test this, we examined animals carrying a mutation in the gene *him-4*, in which linker cell migration is defective, and the cell ends up near the head^19^. In these animals, linker cell death occurs fairly reliably, as determined by onset of nuclear crenellation^19^. However, while engulfment and degradation by neighboring cells can occur, it is inefficient (62 ± 6.9% remaining corpses in 24h adult males, N=50, p<0.0001 compared to wild type, Fisher’s exact test). Thus, U.l/rp cells exhibit specialization for linker cell clearance.

### Apoptotic Engulfment Genes Are Not Required for Linker Cell Clearance

It was previously reported that linker cell engulfment can occur in animals lacking apoptosis engulfment genes^19^; however, whether the cell is eventually degraded in these settings was not examined. To look at this in detail, we followed linker cell engulfment in animals lacking components of each or both parallel pathways that together drive apoptotic cell engulfment. Animals were scored 24h after the L4-to-adult transition, when the majority of animals have cleared the linker cell corpse^24^. We found that linker cell engulfment is only weakly perturbed by mutations in some known engulfment genes (Table S2). Furthermore, double mutants between engulfment genes in different pathways do not strongly affect linker cell clearance. Consistent with this observation, when we expressed MFG-E8-GFP, a phosphatidylserine binding protein^26,27^, in U cell descendants, we did not observe fluorescence accumulation around the linker cell (5 lines, >100 animals scored in total). Nonetheless, we did find that mutations in the phagosome maturation gene *sand-1* significantly inhibit clearance of linker cell debris (Table S2).

Thus, while genes promoting cell corpse degradation following engulfment are shared between linker cell death and apoptosis, upstream genes required for phagocytosis and the initiation of degradation are different. Linker-cell-specific engulfment genes remain, therefore, to be discovered.

### RME-4/DENND1 Promotes Linker Cell Engulfment and Degradation

To identify components of the linker cell engulfment machinery, we sought mutants in which linker cell engulfment and/or degradation is defective. To do so, we mutagenized hermaphrodites carrying a *lag-2*p::GFP linker cell reporter and a *him-5* mutation, which increases the proportion of male progeny (Figure S2A). After two generations, males were screened for linker cell persistence. We previously showed that animals defective in linker cell death exhibit gonadal blockage, and can be sterile^19^. We therefore directly collected sperm from mutant males by impaling a needle into the gonad, and used this sperm to artificially inseminate wild-type hermaphrodites^28^ (Figure S2B-D, see Methods for details). Cross progeny were then used to establish mutant lines.

From this screen, we identified 50 mutant strains with defects in linker cell death and degradation. One mutant, *ns410*, exhibits persistent refractile linker cell corpses, and was further studied (Figure 2A,B). Using standard genetic mapping, coupled with whole genome sequencing, we identified a mutation predicted to generate a G100E alteration in the DENN domain protein RME-4, homologous to vertebrate DENND1^29^ (Figure S3A). Linker cell degradation is restored to *ns410* mutants by expressing a fosmid spanning the genomic region of *rme-4* (Figure 2B). Furthermore, two other *rme-4* alleles, *ns412* (S217N; isolated in our screen) and *b1001* (G257D; a previously characterized loss-of-function lesion^29^) also promote linker cell degradation defects, as does the *rme-4*(*tm1865*) deletion allele (Figure S3B). *rme-4*(*tm1865*) mutants exhibit a weaker linker cell survival defect, perhaps because the truncated RME-4 protein produced in this mutant is present at half wild-type levels^29^. RNAi against *rme-4* exacerbates the defect of animals carrying this allele (Figure S3B). Taken together, these results demonstrate that the *rme-4* gene is required for linker cell corpse degradation.

**Figure 2.**
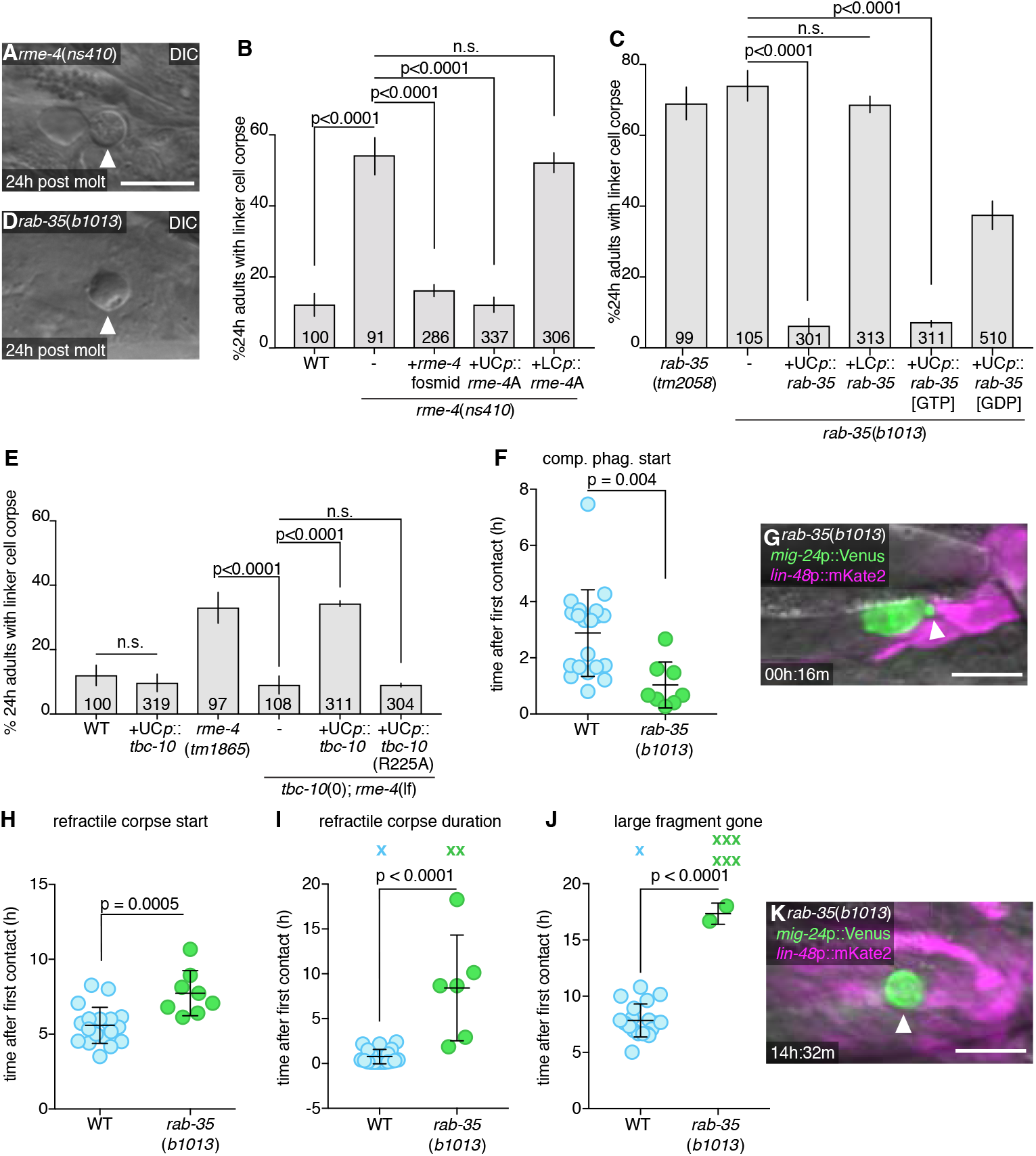
RAB-35, RME-4 (DENND1), and TBC-10 (TBC10D1A) Regulate Linker Cell Clearance. (A) DIC image of persistent linker cell corpse (arrowhead) in *rme-4*(*ns410*). Scale bar, 10 μm. (B,C) Linker cell degradation in indicated genotypes. Strains contain *lag-2*p::GFP linker cell reporter and *him-5*(*e1490*). UCp, *lin-48*p. LCp, *mig-24*p. Number of animals scored inside bars. Average of at least three independent lines. Error bars, standard error of the proportion or standard error of the mean. n.s., p>0.05, Fisher’s exact test. (D) DIC image of persistent linker cell corpse (arrowhead) in *rab-35*(*b1013*). (E) Histogram details as in B. (F) Competitive phagocytosis onset in individual animals (circles). Bars, mean ± sd. Student’s t-test. WT, wild-type. (G) Arrowhead, early competitive phagocytosis in *rab-35*(*b1013*). Details as in Figure 1. (H-J) Quantification of linker cell events as in (F). X, Event persisting or not observed by end of imaging; not included in statistical analysis. (K) Arrowhead, persistent large cell fragment in *rab-35*(*b1013*). Strain and image details as in G.

RME-4 is widely expressed and found in the linker cell and in its engulfing cells (Figure S3C). To determine where RME-4 functions, we expressed cDNAs for either the *rme-4*A or *rme-4*B transcript (Figure S3A) in either the linker cell or the engulfing cells. We found that the longer A isoform restores linker cell degradation when expressed in U cell descendants, but not when expressed in the linker cell (Figure 2B). B isoform expression in the engulfing cells has no effect (Figure S3B). Thus, RME-4A is the active isoform driving linker cell engulfment, and it does so by acting in the engulfing cells.

### RAB-35 GTPase Is a Key Linker Cell Degradation Regulator Controlled by RME-4/DENND1 and TBC-10/TBC1D10A

The RME-4 DENN domain was previously proposed to drive the guanine nucleotide exchange reaction (GEF function) of the small GTPase RAB-3 5^29,30^. The *ns410, ns412*, and *b1001 rme-4* alleles are predicted to cause single amino-acid changes within this domain, suggesting that RME-4 could function as GEF during linker cell degradation. To test this directly, we examined two *rab-35* mutants, *tm2058* and *b1013*, that carry a small deletion surrounding the start codon, and an early Q69Ochre stop mutation, respectively (Figure S3D). We found that both lesions inhibit linker cell corpse degradation, and persistent corpses resemble those seen in *rme-4* mutants (Figure 2C,D). While *rab-35* is broadly expressed (Figure S3E), we found that linker cell degradation is restored to *rab-35* mutants by expression of *rab-35* cDNA in engulfing cells, but not in the linker cell (Figure 2C). Thus, RAB-35 is required in engulfing cells for linker cell degradation.

RAB-35 and other small GTPases cycle between GTP and GDP bound forms. If RME-4 functions as a RAB-35 GEF, we expect RAB-35[GTP] to be the active form of the protein in linker cell degradation. Indeed, we found that expression of RAB-35(Q69L), predicted to lock the protein in the GTP-bound configuration, fully restores linker cell degradation to *rab-35*(*b1013*) mutants (Figure 2C). A GDP-bound mimetic, RAB-35(S24N), however, only partially rescues *rab-35*(*b1013*) linker cell defects. Furthermore, we found that RAB-35(S24N) protein selectively binds RME-4A (but not RME-4B) in a yeast two-hybrid assay^29^ (Figure S3F). Thus, RAB-35 functions with RME-4 in engulfing cells, and RAB-35[GTP] is likely the relevant active form driving linker cell clearance.

We next sought to identify the GTPase activating protein (GAP) completing the RAB-35 GTPase cycle. We reasoned as follows: weak RME-4 (GEF) mutations should result in some RAB-35 [GTP] production, but perhaps not enough for efficient linker cell degradation. Combining an *rme-4*(weak) mutation with a RAB-35 GAP mutation, which would block RAB-35 [GTP] hydrolysis, could, however, allow sufficient accumulation of RAB-35[GTP] for efficient linker cell degradation. We therefore screened for the effects of mutations in *C. elegans* homologs of the known mammalian Rab35 GAP genes *tbc-7/Tbc1D24, tbc-10/Tbc1D10A*, and *tbc-13/Tbc1D13*^31^. Mutations or knockdown of these genes alone have no effect on linker cell corpse degradation, nor do they rescue linker cell degradation defects of animals carrying the strong *rme-4*(*ns410*) mutation, in which no RAB-35[GTP] is predicted to accumulate (Figure S3G). However, combining the *tbc-10*(*gk388086*) allele with the weak *rme-4*(*tm1865*) allele restores normal linker cell degradation (Figure 2E). Thus, TBC-10 is likely the RAB-35 GAP.

Supporting this notion, while TBC-10 protein is widely expressed (Figure S3H), specific expression of TBC-10 in U cell descendants of *rme-4*(*tm1865*); *tbc-10*(*gk388086*) double mutants reintroduces the linker cell degradation defect. Similar expression of a TBC-10(R225A) protein, containing a lesion in the putative catalytic arginine finger, has no effect (Figure 2E).

Taken together, our results suggest that RAB-35 [GTP] is a key regulator of linker cell degradation, and that its activity is controlled by RME-4/GEF and TBC-10/GAP.

### RAB-35 Promotes Timely Onset of Linker Cell Engulfment and Is Required For Subsequent Phagosome Maturation

To define more specifically the defects associated with loss of RAB-35, we imaged *rab-35*(*b1013*) mutants using the microfluidic setup, and quantified hallmark linker cell death and degradation events over > 24h (Movie S2; Table S1). We found no significant differences compared to wild-type animals in the onset or duration of developmental milestones (e.g. tail-tip retraction or appearance of rays; Figure S4A-C), or in appearance of linker cell death hallmarks (e.g. nuclear crenellation; Figure S4D-I). However, competitive phagocytosis of the linker cell begins prematurely in *rab-35* mutants (Figure 2F), initiating before the linker cell has the opportunity to intercalate between the U.l/rp cells (Figure 2G). While subsequent linker cell splitting occurs at the same time as in wild-type animals (Figure S4G), formation of the larger refractile corpse is delayed (Figure 2H), and refractility can last much longer (Figure 2I). Mutations in *rab-35* delay degradation of the large fragment, if it occurs at all (Figure 2J,K). The smaller linker cell fragment is degraded as in the wild type (Figure S4H,I). Thus, RAB-35 plays roles in both engulfment initiation and phagosome maturation.

Consistent with a role in engulfment initiation, we found that RAB-35, tagged with the fluorescent reporter YFP, localizes to extending pseudopods early on, and remains enriched around the larger linker cell-containing phagosome for an extended period of time, until late stages of linker cell degradation. (Figure 3A-C).

**Figure 3.**
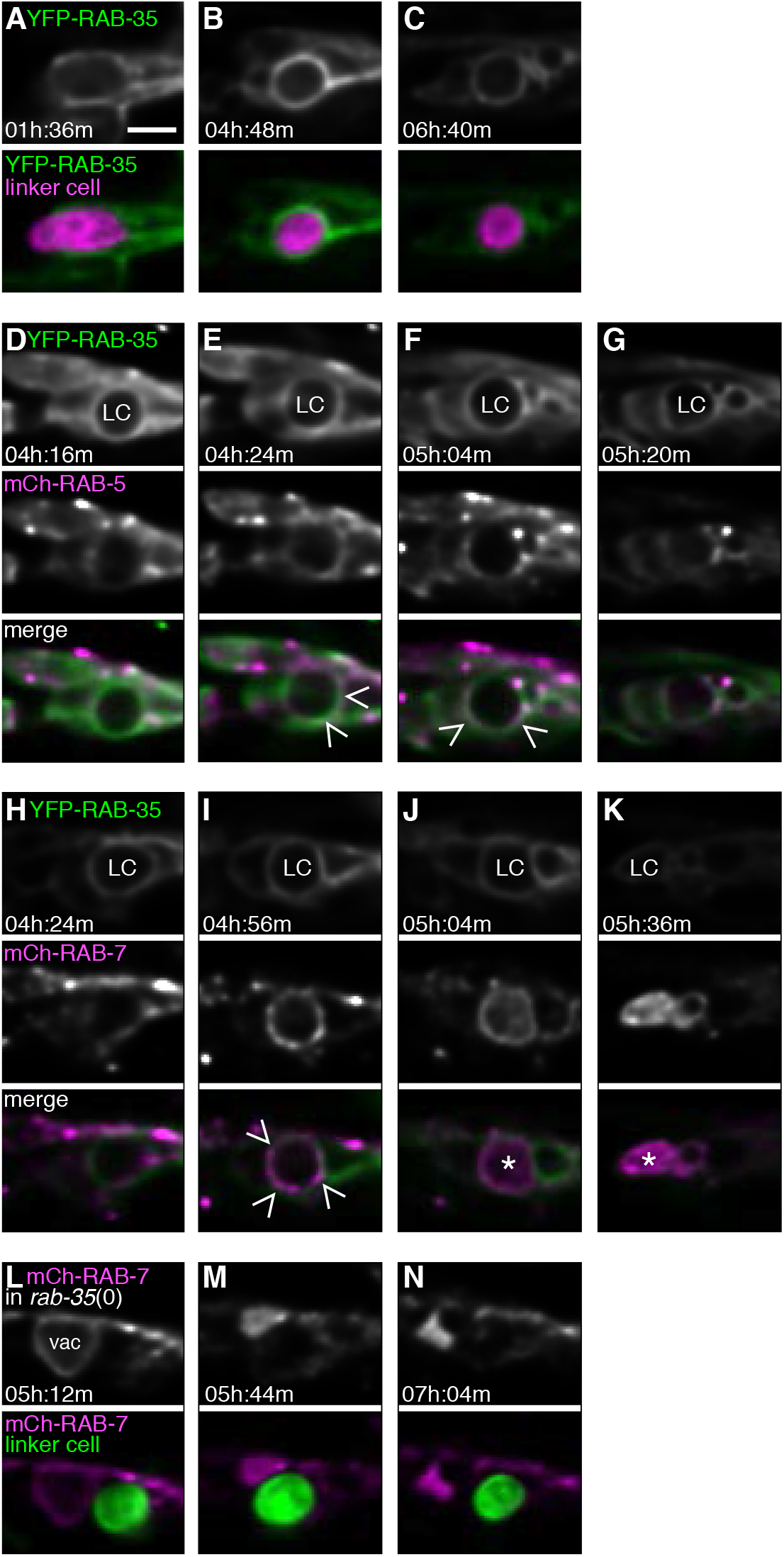
RAB-35 Colocalizes with RAB-5 and RAB-7 and Is Required for Their Recruitment around the Linker Cell. (A-C) Top: Localization of YFP-RAB-35 within U.l/rp cells. Bottom: YFP-RAB-35 (green), and linker cell (*mig-24*p::mKate2; magenta). Scale bar, 5 μm. (D-G) Same format as (A), except mCherry-RAB-5 imaged, and linker cell not fluorescently labeled. Caret, site of colocalization. LC, linker cell. (H-K) Same format as (D), except mCherry-RAB-7 also imaged. Caret, site of colocalization. Asterisk, accumulation within phagosome. (L-N) mCherry-RAB-7 fails to surround linker cell in *rab-35*(*b1013*) mutant. Same format as above.

Some aspects of phagosome maturation have been previously characterized. We therefore sought to address where in this process RAB-35 functions. To do so, we first generated animals expressing both YFP-RAB-35 and either mCherry-RAB-5, marking early phagosomes, mCherry-RAB-7, marking late phagosomes, or CTNS-1-mKate2, marking the phagolysosome (Figures 3A-K; S5A-C; Table S1). In wild-type animals, RAB-35 and RAB-5 transiently co-localized on the phagosome membrane in 5/6 animals examined (Figure 3E,F). In three of six animals, RAB-5 enrichment was seen in only a single frame over a 20h experiment (images acquired every 8 min), suggesting that RAB-5 can move quickly on and off the phagosome membrane. RAB-7 and RAB-35 co-localization is more sustained (Figure 3I, 7/7 animals). Furthermore, as RAB-35 disappears from the phagosome membrane, mCherry-RAB-7 fluorescence accumulates within the phagosome (Figure 3J,K). Such internalization of RAB-7 had not been previously reported in other settings. CTNS-1 and RAB-35 exhibit minimal co-localization (Figure S5A-C), suggesting that RAB-35 is removed from the phagosome membrane once lysosomal fusion occurs.

We next examined reporter localization in *rab-35*(*b1013*) mutants. We found that RAB-5 fails to localize to the phagosome membrane in these animals (Figure S5D-F, 5/7 animals). Similarly, RAB-7 does not accumulate on the phagosome surface, and is not taken up into the phagosome (Figure 3L-N, 5/5 animals).

Our studies, therefore, support the idea that RAB-35 acts early in phagosome maturation, and is required for recruiting RAB-5 and RAB-7 for proper degradation to ensue.

### The Small GTPase ARF-6 Blocks Linker Cell Clearance

To delineate the molecular mechanism by which RAB-35 controls linker cell engulfment onset and degradation, we pursued studies of another mutant, *ns388*, isolated in our genetic screen. Animals carrying this lesion have similar linker cell defects to *rab-35* mutants (Figure 4A,B), but do not harbor mutations in *rab-35, tbc-10*, or *rme-4*.

**Figure 4.**
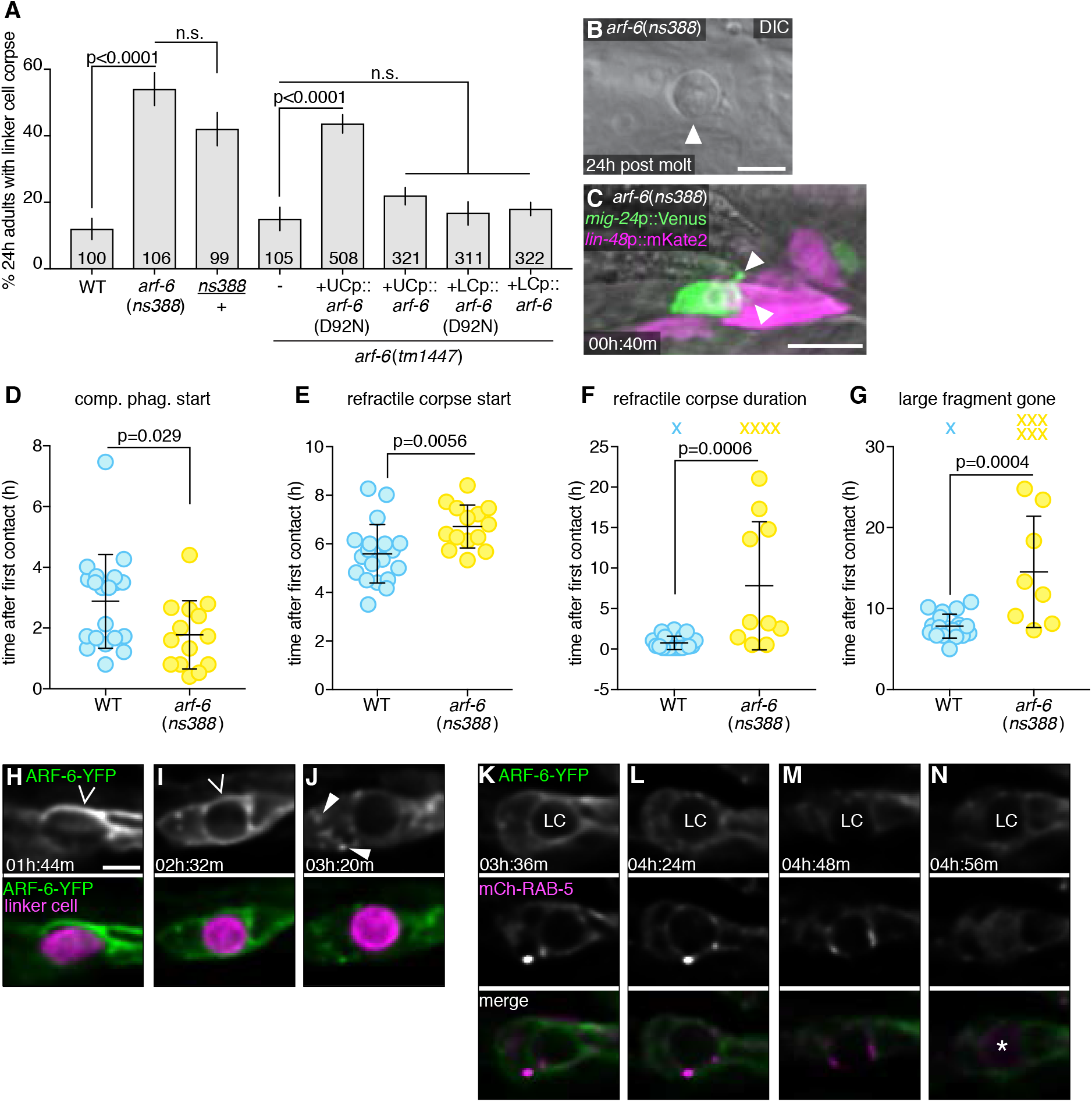
ARF-6 Inhibits Linker Cell Degradation. (A) Linker cell degradation in indicated genotypes. Details as in Figure 2B. (B) DIC image of persistent linker cell corpse (arrowhead) in *arf-6*(*ns388*) male. Scale bar, 5 μm. (C) Arrowheads, premature competitive phagocytosis in *arf-6*(*ns388*). Scale bar, 10 μm. (D-G) Timing of linker cell clearance events in *arf-6*(*ns388*). Quantification as in Figure 2F,H-J. (H-N) Localization of ARF-6-YFP and phagosome maturation markers. Scale bar, 5 μm. (H-J) Top: Localization of ARF-6-YFP within the U.l/rp cells. Caret, ARF-6 on phagosome membrane. Arrowhead, ARF-6 in intracellular puncta. Bottom: ARF-6-YFP (green), and linker cell (*mig-24*p::mKate2; magenta). (K-N) Localization of ARF-6-YFP (top) and mCherry-RAB-5 (middle). LC, linker cell. Asterisk, accumulation of RAB-5 within phagosome.

Using whole genome sequencing and standard genetic mapping, we found that *ns388* animals contain a point mutation predicted to cause a D92N mutation in the small GTPase ARF-6 (Figure S6A). Importantly, and unlike *rab-35* mutations, a single *arf-6*(*ns388*) allele is sufficient to block linker cell degradation. Thus, *arf-6*(*ns388*) is a dominant allele (Figure 4A). The *arf-6*(*tm1447*) putative null allele, which lacks most of the coding region, and has no ARF-6 expression by Western blot^32^, exhibits normal linker cell clearance (Figure 4A), suggesting that *arf-6*(*ns388*) is a gain-of-function and not a dominant-negative allele. To confirm that *arf-6* is indeed the relevant gene, we generated two independent CRISPR alleles, *ns751* and *ns752*, recreating the *arf-6*(*ns388*) lesion. Both promote linker cell defects similar to those of *arf-6*(*ns388*) mutants (Figure S6B). Furthermore, while *arf-6* is widely expressed (Figure S6C-D), expression of an *arf-6*(*ns388*) cDNA in engulfing cells blocks linker cell clearance (Figure 4A). Wild-type *arf-6* expression in engulfing cells does not affect linker cell death, nor does expression of either *arf-6*(*ns388*) or *arf-6*(+) cDNAs in the linker cell (Figure 4A).

Our results, therefore demonstrate that gain of ARF-6 function blocks linker cell clearance, and suggest that ARF-6 normally functions as a linker cell clearance inhibitor.

### ARF-6(gf) Promotes Premature Competitive Phagocytosis Onset and Delays Linker Cell Degradation

To understand how ARF-6(gf) interferes with linker cell clearance, we imaged *arf-6*(*ns388*) animals in the microfluidic device and followed hallmark linker cell death and degradation events (Movie S3; Table S1). Except for a slight advance in cuticle shedding, no defects in animal development or linker cell killing are seen in *arf-6*(*ns388*) mutants (Figure S4). However, as with *rab-35* mutants, premature onset of competitive phagocytosis is observed (Figure 4C,D), formation of a refractile linker cell corpse is delayed (Figure 4E), and the period of linker cell refractility is prolonged (Figure 4F). Degradation of the large linker cell fragment is greatly delayed, if it occurs at all (Figure 4G).

The similar defects exhibited by *arf-6*(*ns388*) and *rab-35* mutants suggests that the proteins encoded by these genes may localize in a similar manner. To test this idea, we expressed ARF-6-YFP in the U.l/rp cells and followed its localization changes during linker cell death and degradation (Figure 4H-M). ARF-6-YFP localizes to U.l/rp pseudopods (Figure 4H; Table S1), remains enriched on the phagosome membrane after cell splitting (Figure 4I), and then rapidly translocates to intracellular puncta (Figure 4J). Importantly, enrichment of mCherry-RAB-5 around the phagosome occurs after ARF-6-YFP removal (Figure 4K-M, 5/6 animals). As with RAB-7, we also observed RAB-5 accumulation in the phagosome interior (Figure 4N, 6/6 animals).

Thus, ARF-6 and RAB-35 localize to similar structures surrounding the dying linker cell. However, while RAB-35 normally promotes linker cell degradation, ARF-6 blocks this process.

### ARF-6 Function Is Regulated by CNT-1/ACAP2 and EFA-6/EFA6

To understand the dominant nature of the ARF-6(D92N) lesion, we attempted to assess the consequences of locking ARF-6 protein in the GTP or GDP bound states^33^. We used CRISPR to generate *arf-6* mutants encoding T44N and Q67L mutations (Figure S6A), predicted to accumulate ARF-6[GDP] or ARF-6[GTP] proteins, respectively. Neither mutant, however, has linker cell defects, perhaps because they destabilize the protein. Consistent with this interpretation, the *arf-6*(Q67L) allele behaves as a null in genetic assays (see below). We did however, serendipitously isolate from our CRISPR mutagenesis an *arf-6*(I42M,P43T) double mutant with linker cell clearance defects. Based on an Arf6:Arf6GAP co-crystal protein structure^34^, these lesions may interfere with GAP binding to ARF-6, and subsequent GTP hydrolysis. D92 of Arf6 forms intramolecular hydrogen bonds with R95 (Figure 5A). In the D92N mutant, these hydrogen bonds are lost (Figure 5B), which may result in loss of stability of adjacent interactions. Both P43 and I42 of Arf6 form hydrophobic interactions with W451 and I462 on the ArfGAP (Figure 5C). These interactions are lost in the P43T;I42M mutations (Figure 5D). Therefore, ARF-6[GTP] appears to inhibit phagosome maturation during linker cell clearance, and the mutants we isolated may hinder GAP binding.

**Figure 5.**
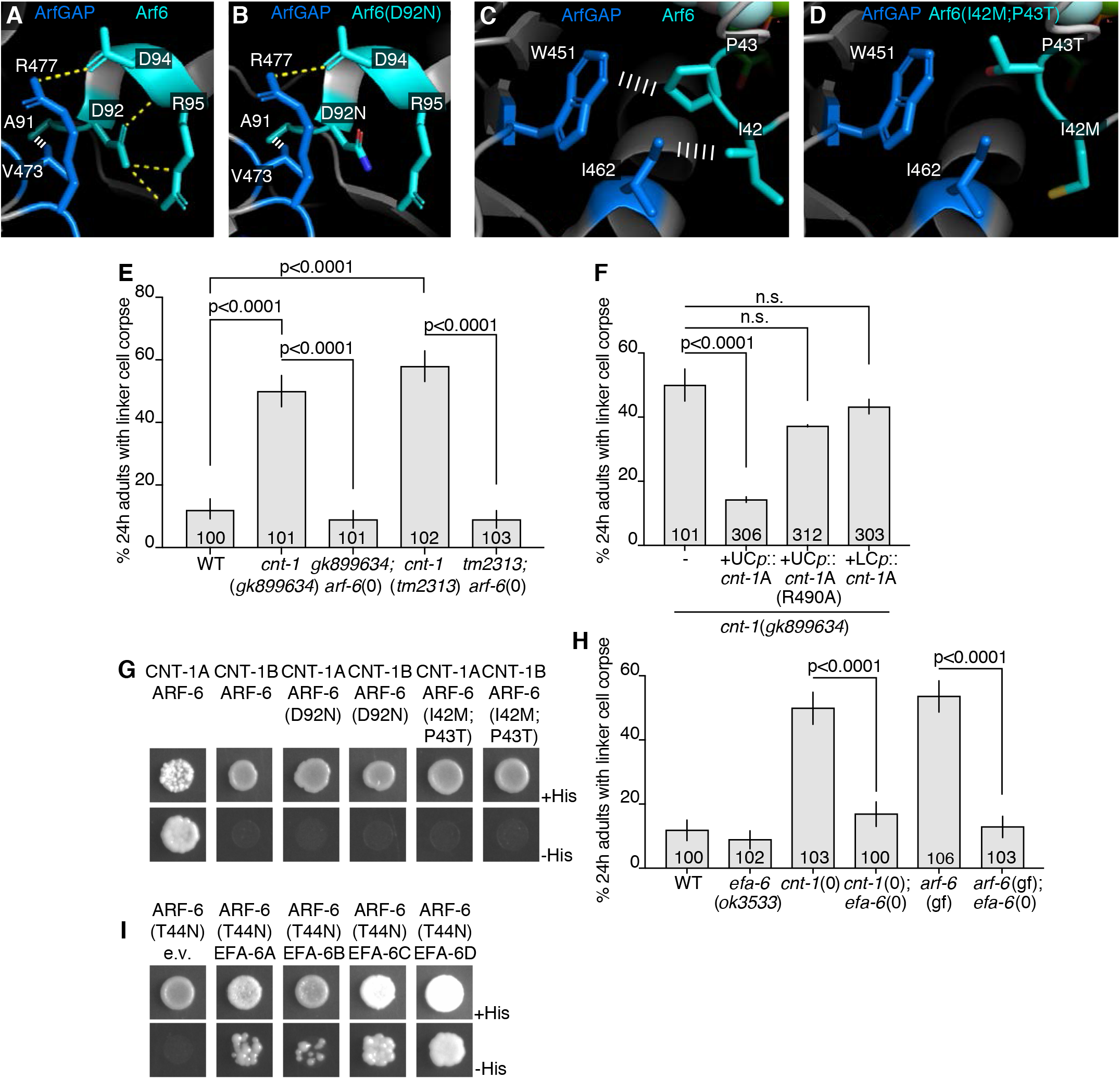
EFA-6 (EFA6) and CNT-1 (ACAP2) Regulate ARF-6. (A) Protein structures of human Arf6 (teal) and ArfGAP (blue) from^35^. PDB: 3LVQ. Relevant amino acids indicated in white. Yellow dotted line, hydrogen bonds. White ladder, hydrophobic interactions. Amino acids shown are conserved in *C. elegans* ARF-6 and CNT-1. (B) D92N mutation mapped onto Arf6. Protein structure information as in (A). (C) Different region of Arf6 and ArfGAP. Protein structure information as in (A). (D) I42M, P43T mutations mapped onto Arf6. (E-F) Histogram details as in Figure 2B. (G) Yeast two-hybrid assay with LexA-CNT-1A or LexA-CNT-1B as bait, GAD-ARF-6, GAD-ARF-6(D92N) or GAD-ARF-6(I42M,P43T) as prey. Top: histidine present. Bottom: histidine absence. Growth on -His plates indicates physical interaction. (H) Histogram details as in Figure E. (I) As in (G), with LexA-ARF-6[T44N] (GDP) as bait, GAD-EFA-6A-D as prey.

To test this idea further, we reasoned that mutations in a relevant ARF-6 GAP would exhibit linker cell clearance defects, as they would lock ARF-6 in the GTP-bound state. From a screen of candidate mutations (Figure 5E; Table S3), we found that two independent putative null alleles of *cnt-1*, encoding a ubiquitously-expressed protein similar to vertebrate Acap2, block linker cell clearance (Figures 5E; S6E,F). Expression of the CNT-1A isoform in U.l/rp cells, but not the linker cell, fully rescues linker cell defects of *cnt-1* mutants (Figure 5F); whereas CNT-1B expression gives partial rescue (Figure S6G). Rescue is abolished by mutating the catalytic arginine of CNT-1A (Figure 5F). Importantly, ARF-6 is required for the linker cell defects of *cnt-1* mutants, as an *arf-6*(*tm1447*) loss-of-function mutation restores linker cell clearance to *cnt-1* mutants (Figure 5E). Consistent with the idea that the D92N and I42M;P43T mutations disrupt GAP binding, we found that while wild-type ARF-6 interacts with CNT-1A (but not CNT-1B) in a yeast two-hybrid assay; both ARF-6(D92N) and ARF-6(I42M;P43T) fail to interact with CNT-1A (Figure 5G). Thus, ARF-6[GTP] is likely the active form of the protein blocking linker cell removal, and CNT-1 is the ARF-6 GAP for linker cell clearance.

Similar reasoning used to identify TBC-10 as a RAB-35 GAP allowed us to identify EFA-6, homologous to the mammalian Arf6 GEF Efa6^35^, as the putative ARF-6 GEF for linker cell clearance. While the *efa-6*(*ok3533*) deletion allele has no linker cell degradation defect, this allele restores linker cell clearance to mutants carrying null alleles of *cnt-1* (Figure 5H; *cnt-1* (*tm2313*); *efa-6*(*ok3533*): 14 ± 3.4%, N = 102, p<0.0001 compared to *cnt-1*(*tm2313*), Fisher’s exact test). Furthermore, *efa-6*(*ok3533*) also blocks the dominant defects of *arf-6*(*ns388*) and *arf-6*(*ns763*[I42M;P43T]) mutants, strengthening our assessment that ARF-6 GTP loading is required for the function of these ARF-6 gain-of-function proteins (Figures 5H; S6H). While EFA-6 is expressed in the linker cell and U.l/rp cells (Figure S6I), expression of EFA-6 isoforms in the U.l/rp cells partially restores linker cell clearance defects to *efa-6*(*ok3533*); *arf-6*(*ns388*) double mutants (Figure S6H). Furthermore, EFA-6 isoforms interact with ARF-6[GDP] in a yeast two-hybrid assay (Figure 5I).

Taken together, our results suggest that ARF-6 inhibits linker cell phagocytosis onset, and blocks linker cell degradation together with its regulators EFA-6/GEF and CNT-1A/GAP.

### RAB-35 Binds CNT-1A and Drives ARF-6 Removal From Phagosome Membranes

*rab-35*(0) and *arf-6*(gf) mutants exhibit similar linker cell clearance defects. We wondered, therefore, whether RAB-35 and ARF-6 proteins act in the same pathway. We found that putative null mutations in *arf-6* or *efa-6* almost fully restore linker cell clearance to *rab-35* mutants (Figure 6A; S7A). The *arf-6*[Q67L] CRISPR mutant also suppresses *rab-35*(0) defects (Figure S7B), confirming this allele inactivates ARF-6 and is not an ARF-6[GTP] mimetic (see above). Linker cell clearance defects are restored to *rab-35*(*b1013*); *arf-6*(*tm1447*) double mutants by expressing wild-type *arf-6* in U.l/rp cells, but not in the linker cell (Figure S7C). Furthermore, neither mutations in *cnt-1* nor the dominant *arf-6* allele enhance the linker cell defects of *rab-35* mutants (Figure S7D). Finally, overexpression of YFP-RAB-35 in engulfing cells restores linker cell clearance to *arf-6*(*ns388*) mutants (Figure 6A). Taken together, these results suggest that RAB-35 normally functions to inactivate ARF-6.

**Figure 6.**
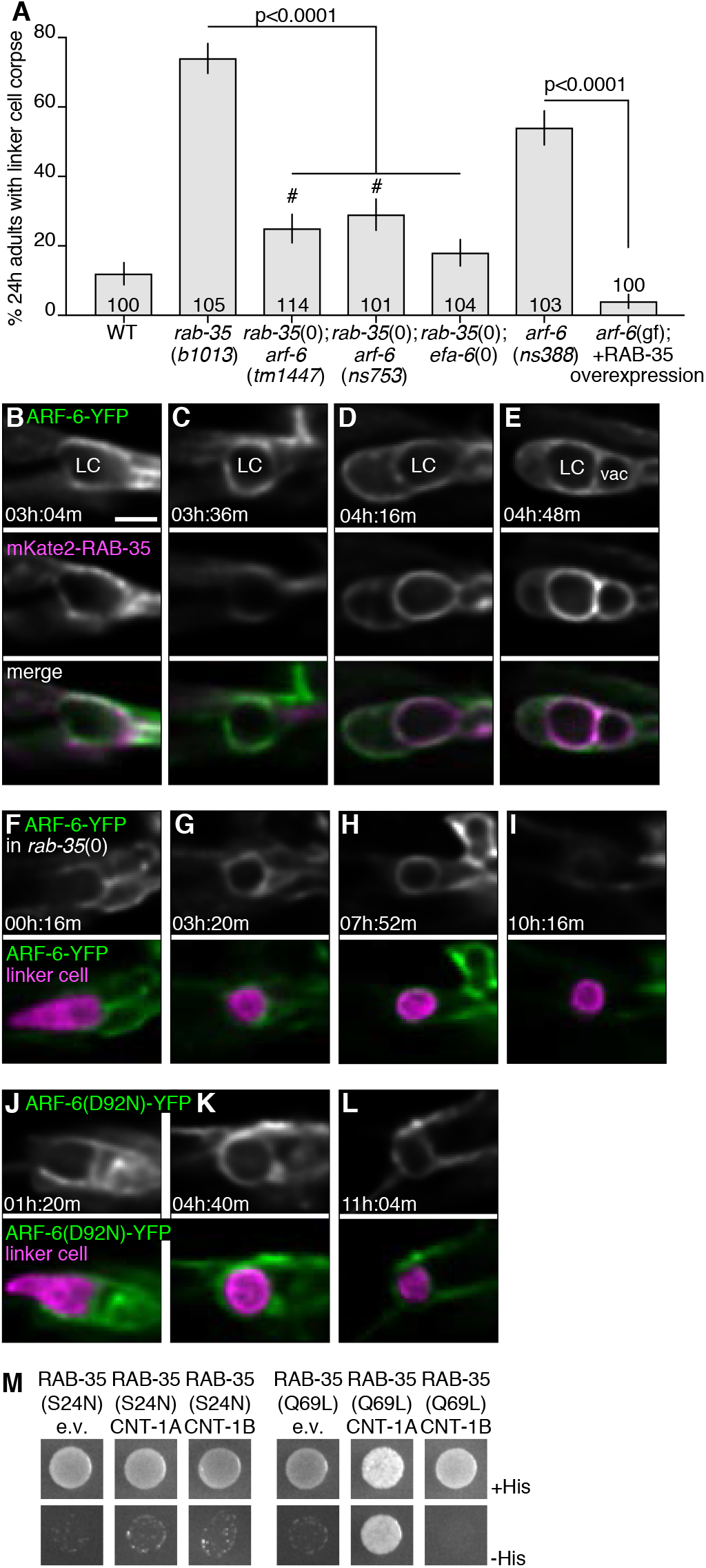
RAB-35 Removes ARF-6 from Phagolysosome Membranes. (A) Histogram details as in Figure 2B. #, significant difference compared to *rab-35*(*b1013*) and wild type. (B-L) Localization of indicated proteins with details as in Figure 3. (F-I) In *rab-35*(*b1013*) mutant. (M) Yeast two-hybrid assay with LexA-RAB-35[S24N] (GDP) or LexA-RAB-35[Q67L] (GTP) as bait, Gal4-AD (GAD)empty vector, GAD-CNT-1A, or GAD-CNT-1B as prey. Details as in Figure 5.

To understand how RAB-35 inhibits ARF-6, we first imaged mKate2-RAB-35 and ARF-6-YFP localization in the same animal. Consistent with our other imaging studies, the two proteins initially co-localize to extending U.l/rp cell pseudopods (Figure 6B). RAB-35 then accumulates on the phagosome membrane, while ARF-6 enrichment fades (Figure 6C-E, 8/9 animals). Next, we examined ARF-6-YFP dynamics in *rab-35*(*b1013*) animals (Figure 6F-I). Remarkably, we found that ARF-6-YFP remains on the membrane longer than in wild type animals (Figure 6G,H, 5/6 animals; ARF-6-YFP removed 4.6 ± 0.9h after first contact in wild type animals vs 15.4 ± 9.7h in *rab-35*(*b1013*) where the corpse persists, p=0.023, Student’s t-test). Furthermore, in wild-type animals, ARF-6(D92N) remains enriched on phagosome membranes longer than ARF-6-YFP (Figure 6J-L, 4/4 animals; 19.1 ± 11.4h after first contact, p=0.01 compared to WT, Student’s t-test).

These results raise the possibility that RAB-35 promotes linker cell clearance, at least in part, by facilitating ARF-6 removal from the phagosome. Consistent with this notion, an ARF-6(D92N) gain-of-function protein is more effective in blocking linker cell clearance than ARF-6(D92N) lacking a myristoyl group addition sequence, required for membrane localization^36^ (Figure S7C,E).

To determine whether ARF-6 localization is regulated by direct binding to RAB-35, we assessed their interactions in a yeast two-hybrid system (see Methods). While we did not detect any interactions between these two proteins, we found that RAB-35[GTP], the active form of RAB-35, binds CNT-1A (and not CNT-1B) (Figure 6M). Thus, RAB-35 may recruit CNT-1A to phagosome membranes, catalyzing conversion of active ARF-6[GTP] to inactive ARF-6[GDP].

### The RAB-35/ARF-6 Module Is Not Required For Apoptotic Cell Clearance

Given the roles of RAB-35 and ARF-6 in the clearance of a cell dying by LCD, we wondered whether these proteins play similar roles in apoptotic cell clearance. To test this, we counted the number of persisting apoptotic cells in 3-fold embryos, after nearly all developmental cell death has taken place. In *him-5* mutant control animals, we counted 0.10 ± 0.30 persisting refractile corpses on average (N=21). In *arf-6*(*ns388*) mutants, 0.47 ± 0.60 corpses were observed (N=21, p=0.0131, Student’s t-test). *rab-35*(*b1013*) mutants exhibited 0.81 ± 0.92 corpses, on average (N=21, p=0.0018, Student’s t-test). These small defects pale in comparison to defects exhibited by mutants defective in canonical apoptotic engulfment genes (e.g. *ced-7; ced-10* double mutant: 16.8 ± 4.2 corpses, N = 11). Furthermore, *rme-4* mutant embryos we tested have no clearance defects (*ns410*: 0.14 ± 0.36 apoptotic corpses, N=14; *ns412*: 0.13 ± 0.33 apoptotic corpses, N=24). We also examined whether *rab-35* mutations exhibit genetic interactions with canonical apoptosis engulfment genes in linker cell death, but found no such interactions (Table S4). Our results, therefore, demonstrate that the RAB-35/ARF-6 module plays specific roles in dismantling the linker cell that are not shared with clearance of apoptotic cells in *C. elegans*.

## Discussion

### A New Engulfment and Degradation Pathway for Non-Apoptotic Dying Cells

Caspase-dependent apoptosis does not account for many cell death events that occur during animal development. Indeed, mice homozygous for knockout alleles of key apoptotic genes, including caspase-3, caspase-9, Apaf-1, or Bax and Bak, can survive to adulthood^37–39^, a surprising observation given the prevalence of cell death in murine development. Inactivation of apoptosis genes also only weakly interferes with degenerative disease progression^40^. LCD, a non-apoptotic cell death process, is prevalent in vertebrates^12^ and may account for a significant fraction of cell deaths that take place during development and in disease.

In this study, we demonstrate that the clearance machinery promoting the removal of the linker cell, which dies by LCD, differs from that used to clear apoptotic cells in *C. elegans*. Our studies uncover a protein network promoting linker cell engulfment and degradation (Figure 7A). We find that RAB-35 and ARF-6 play key early roles, controlling the sequential loading of phagosome maturation factors onto the phagosome membrane (Figure 7B). Our data are consistent with a model in which RAB-35[GTP] promotes elimination of dying cells by blocking the activity of an inhibitor of the process, ARF-6[GTP]. Our studies are the first to implicate a RAB-35/ARF-6 module in the engulfment and degradation of dying cells, define in detail multiple accessory factors, identify the relevant targets (e.g. RAB-7) and, importantly, explore the functions of these proteins in vivo in a living multicellular animal.

**Figure 7.**
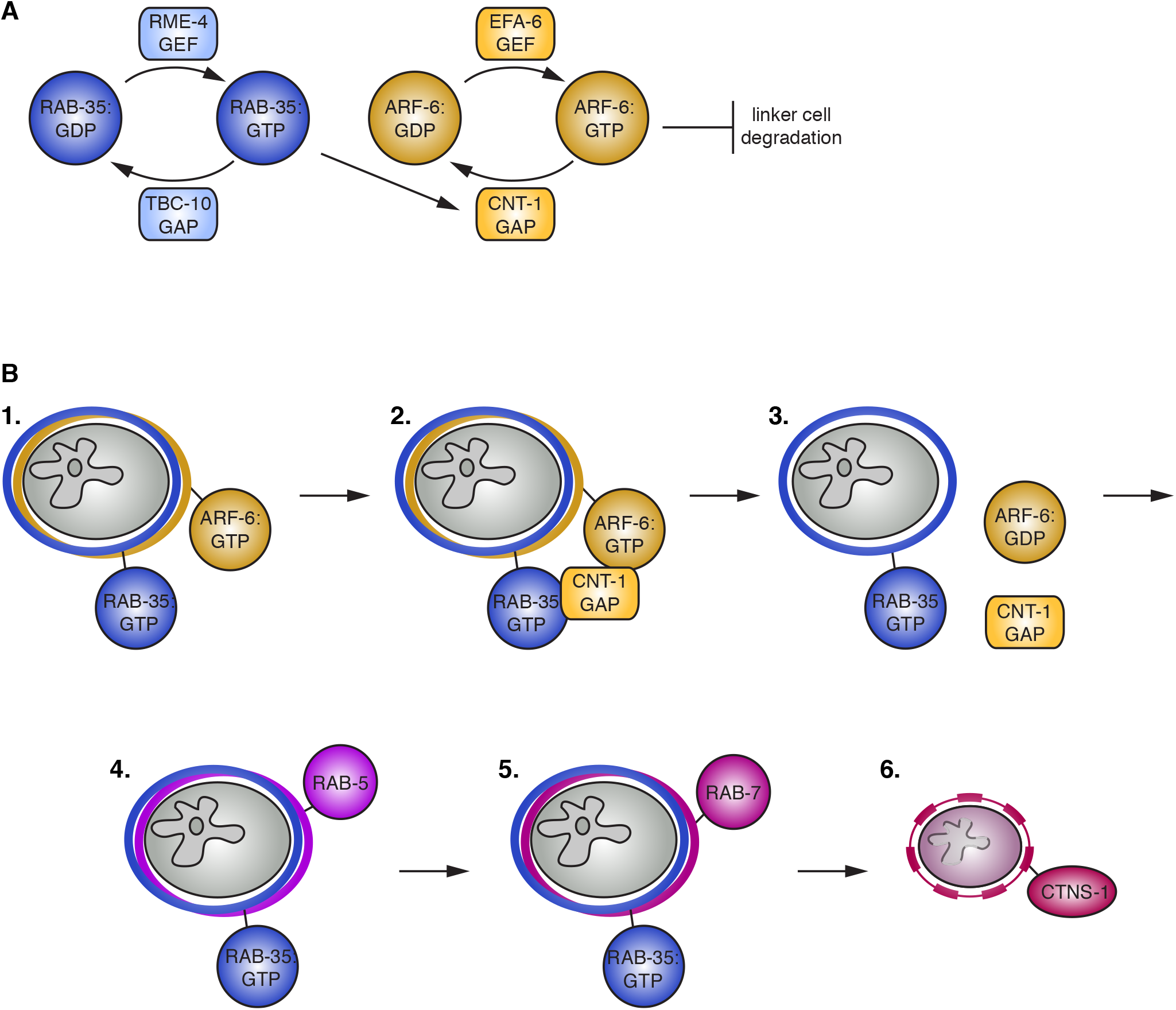
A Model for Control of Linker Cell Degradation by the RAB-35/ARF-6 Module. (A) RAB-35:GTP promotes linker cell degradation by interacting with the ARF-6 GAP CNT-1. CNT-1 is required for turning ARF-6 off and removing it from the membrane. ARF-6:GTP inhibits linker cell degradation. GAP, GTPase activating protein. GEF, guanine nucleotide exchange factor. (B) 1. ARF-6 and RAB-35 surround the nascent phagosome. 2. RAB-35 recruits CNT-1. 3. CNT-1 turns off ARF-6, removing it from the phagosome membrane. RAB-35 remains on the membrane. 4. RAB-5 is recruited. 5. RAB-7 recruitment and RAB-5 loss. 6. RAB-35 is largely gone from the membrane once the lysosome (CTNS-1) fuses with the phagosome.

Although we identify the LCD corpse engulfment pathway in *C. elegans*, a number of studies in other systems raise the possibility that this process may be conserved. For example, in *Drosophila*, nurse cell death is thought to be caspase independent, and knockdown of RAB-35 results in persistent nurse cell nuclei^41^. While the role of RAB-35 was not further investigated, the involvement of this protein in a non-apoptotic setting is intriguing. Furthermore, concerted activities of RAB-35 and ARF-6 in other contexts have been described, including neurite outgrowth, cytokinesis, oligodendrocyte differentiation, maturation of recycling endosomes, and FcγR-mediated uptake of bacteria by macrophage^42–46^. In some of these cases, interactions between RAB-35 and putative ARF-6 GAPs are seen^47^.

### New Ways to Generate ARF[GTP] Mimics

Our genetic analysis of linker cell engulfment yielded important reagents, likely to be of utility in studying ARF-family proteins. We found that a Q67L mutation in ARF-6, which is predicted to mimic the GTP-bound state of the protein, behaves instead as a functional null in our assay. This modification, effective in the context of other GTPases, may, therefore, result in protein instability, or loss of active conformation. However, we found that the ARF-6(D92N) mutant protein, or a protein carrying two lesions, ARF-6(I42M,P43T), appear to mimic ARF-6[GTP], and if anything, enhance protein stability. Based on the published structure of Arf6 binding to the ArfGAP ASAP3^34^, the D92N mutation is adjacent to a residue that coordinates to a key valine in the ArfGAP, whereas the I42M,P43T double mutations modify residues that directly coordinate to known GAP catalysis residues^34^. We propose that these lesions could keep the protein locked in the GTP-bound state, because they prevent GAP protein binding. This interpretation is bolstered by our protein interaction studies. We propose that introducing similar lesions into other ARF proteins, and perhaps into other GTPases, could allow for the development of new GTP-bound mimetics of these proteins.

### Competitive Phagocytosis

In addition to unique molecular components, we found that the mechanics of linker cell engulfment are very different from engulfment of apoptotic cells. Our imaging studies reveal a process of competitive phagocytosis, where two phagocytes compete to engulf portions of the linker cell, resulting in cell splitting. RAB-35 acting through ARF-6 prevents premature activation of this engulfment process. As with the molecular constitutes of linker cell engulfment and degradation, we believe that the process of competitive phagocytosis may also be conserved. In macrophage-less mice, for example, mesenchymal cells engulf dying neighbors during digit formation, and occasionally two mesenchymal cells can be seen engulfing the same dying cell in transmission electron microscopy images^48^. Consistent with this process involving new engulfment regulators, mesenchymal cells do not express the CED-7 homolog ABC1^48^. Similarly, in developing rat cerebellum, more than one non-professional phagocyte can be seen engulfing a dying cell^49^. A related process has been documented during germ cell development in *C. elegans*. Primordial germ cells extend cytoplasm-containing lobes into the adjacent endoderm, and the lobe is excised from the remaining cell, in a process reminiscent of competitive phagocytosis^50^.

The need for engulfment by more than one cell could be related to the size of the dying cell. For example, when macrophages attempt to engulf a particle that is too large, engulfment is stalled^51,52^. Engulfment by two or more cells could solve this problem. However, in this case, cytoplasmic spillage must be avoided. The new engulfment process we describe here accomplishes exactly that.

A process reminiscent of competitive phagocytosis can also be seen when cultured human peripheral blood monocytes engulf a *C. albicans* pathogen^53^. Similarly, cultured mouse monocytes simultaneously engulf a *Leishmania* parasite^54^. One cell engulfs the flagellum, while the other engulfs the posterior pole. In this context, roles for RAB-35 in pathogen clearance^42,47,54^ are especially intriguing, and may suggest underlying mechanisms similar to linker cell clearance.

In summary, we demonstrate that engulfment and degradation of a cell that dies by LCD requires mechanics and machinery different from that used for apoptotic cell engulfment and degradation, and identify a molecular pathway governing this novel form of cell clearance. Given the conservation of LCD, we propose that this ancillary process may be conserved as well.

## Author Contributions

Conceptualization, L.M.K. and S.S.; Methodology, L.M.K., W.K., and S.S.; Software, W.K.; Formal Analysis, L.M.K. and W.K. Investigation, L.M.K. and W.K.; Writing – Original Draft, L.M.K. and S.S.; Writing – Review & Editing, L.M.K., W.K., and S.S.; Funding Acquisition, L.M.K., W.K., and S.S.; Resources, L.M.K., W.K., and S.S.; Visualization, L.M.K., W.K., and S.S.; Supervision, S.S.

## Acknowledgments

We thank Zheng Zhou for sharing unpublished results; Barth Grant, Zheng Zhou, Paul Randazzo, Jacqueline Cherfils, and Shohei Mitani for reagents; Nima Tishbi for technical help; Jordan Ward for the extensive Cel1 protocol; and Shaham Lab members for discussions and critical reading of the manuscript. Some strains were provided by the CGC, which is funded by NIH Office of Research Infrastructure Programs (P40 OD010440). This work was supported by an NIH NRSA Training Grant GM066699 to L.M.K., an HFSP postdoctoral fellowship LT000250/2013-C to W.K., and NIH grants R01NS081490 and R01HD078703 to S.S.

---

**Figure S1.**
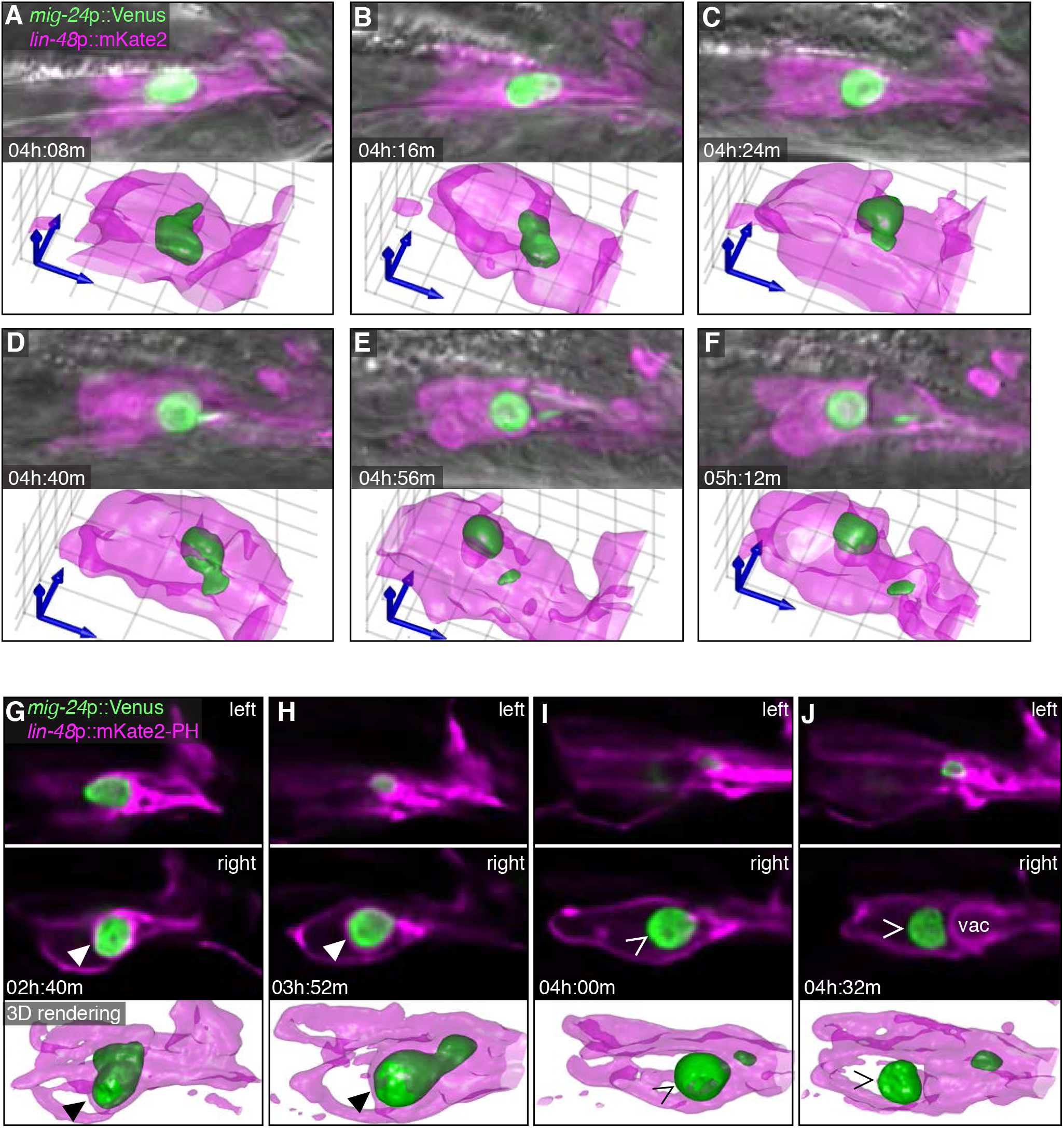
Competitive Phagocytosis Is Dynamic, Related to Figure 1. (A-F) Additional movie frames of animal shown in Figure 1, demonstrating back-and-forth tugging events leading to linker cell splitting. Scale bar, 10 μm. (G-J) Linker cell splitting correlates with loss of mKate2-PH, an open phagosome marker. Scale bar, 10 μm. (G-H) Arrowhead, mKate2-PH plasma membrane marker around linker cell prior to cell splitting (I-J) Caret, mKate2-PH plasma membrane marker absent from phagosome after cell splitting.

**Figure S2.**
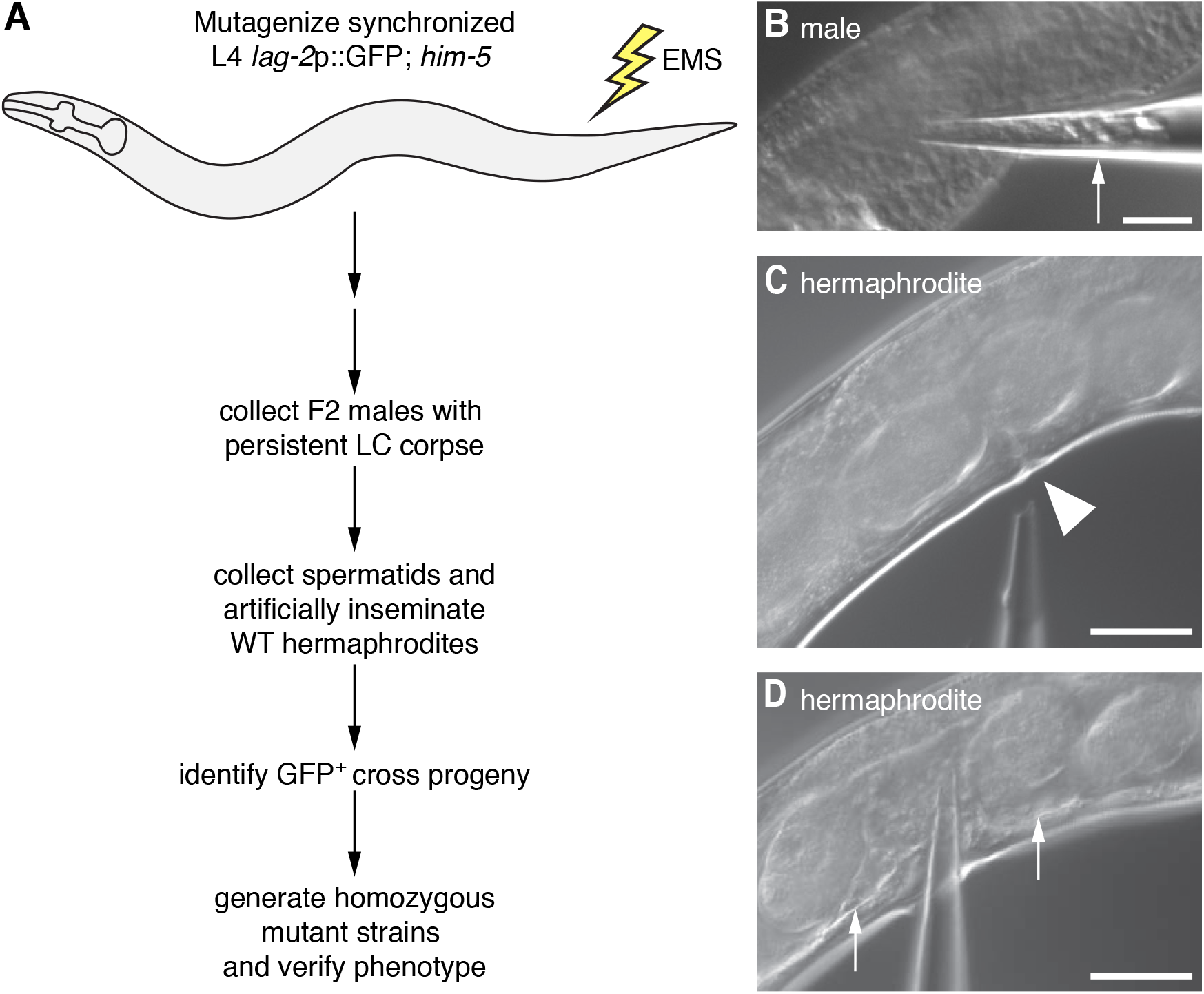
Genetic Screen for Linker Cell Degradation Mutants, Related to Figure 1 and Figure 2. (A) Genetic screen scheme. (B-D) Artificial insemination procedure. (B) Needle filled with buffer plunged into a 24h mutant adult male gonadal cavity. Spermatids automatically flow into the needle (arrow). (C) Needle aligned with a wild type hermaphrodite vulva (arrowhead). (D) Needle inserted into the vulva and spermatids released. Released spermatids fill the hermaphrodite gonadal cavity (arrows).

**Figure S3.**
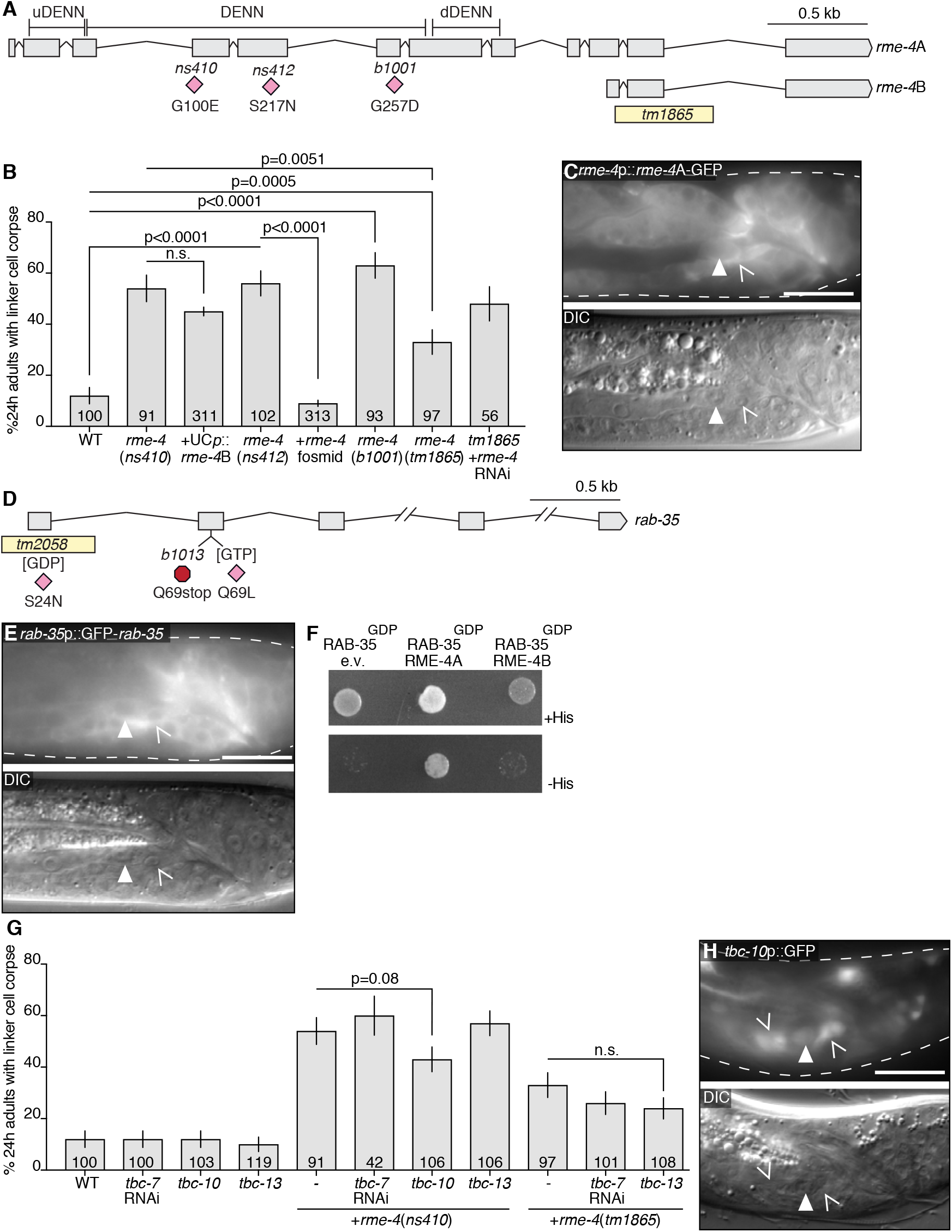
RAB-35, RME-4, and TBC-10 Reagents and Interactions, Related to Figure 2. (A) *rme-4* gene locus. Pink diamonds, locations of point mutations. Yellow box, deletion allele *tm1865*. (B) Histogram details as in Figure 2B. (C) *rme-4* translational reporter expression. Arrowhead, linker cell. Caret, U.lp or U.rp cell. Top: GFP. Bottom: DIC. Scale bar, 10 μm. (D) *rab-35* gene locus. Yellow box, deletion allele *tm2058*. Pink diamonds, point mutations. Red octagon, early stop allele *b1013*. (E) *rab-35* translational reporter expression. Arrowhead, linker cell. Caret, U.lp or U.rp cell. Top: GFP. Bottom: DIC. Scale bar, 10 μm. (F) Yeast two-hybrid assay with LexA-RAB-35[S24N] (GDP) as bait, and Gal4-AD (GAD) empty vector, GAD-RME-4A, or GAD-RME-4B as prey. Top: histidine present. Bottom: histidine absent. Growth on -His plates, physical interaction. (G) Histogram details as in B. (H) *tbc-10* transcriptional reporter. Arrowhead, linker cell. Caret, U.l/rp cells. Top: GFP. Bottom: DIC. Scale bar, 10 μm.

**Figure S4.**
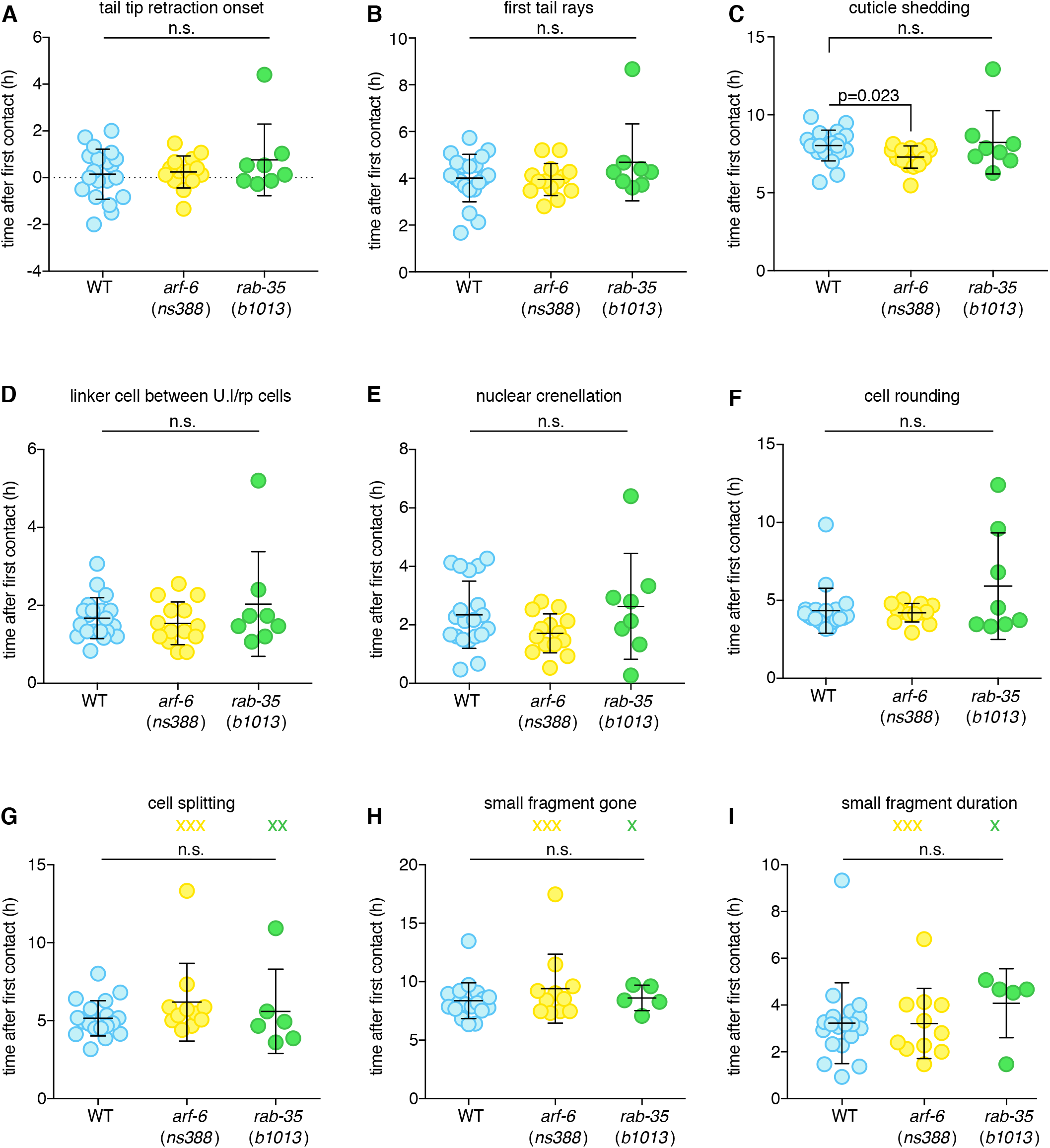
Characterization of Stereotypical Events during Linker Cell Clearance in *rab-35* and *arf-6* Mutants, Related to Figure 2 and Figure 4. (A) (A-I) Strains contain linker cell reporter [*mig-24*p::Venus], U.l/rp cell reporter [*lin-48*p::mKate2], *him-5*(*e1490*). Dots, individual events in single animals in hours with respect to first contact. X, event did not occur; not factored into statistical analysis. Bars, mean ± std. Statistical significance, Student’s t-test.

**Figure S5.**
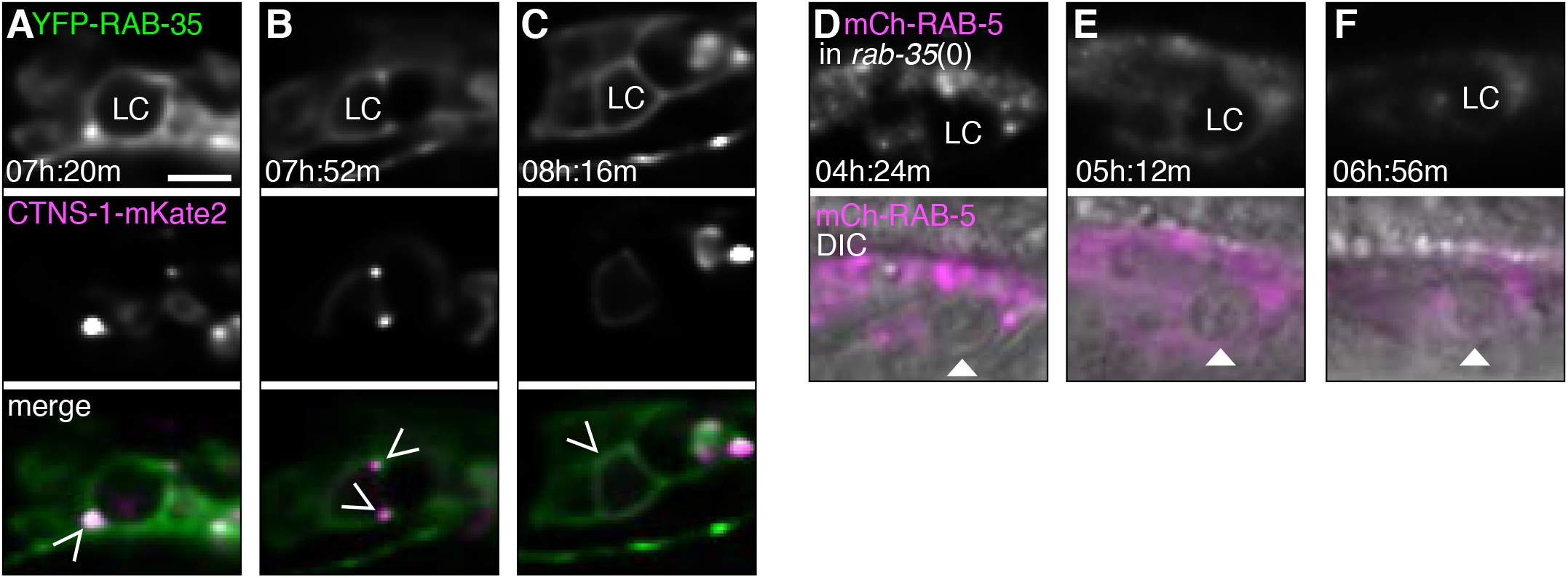
YFP-RAB-35 and CTNS-1-mKate2 Do Not Extensively Colocalize, and RAB-5 Does Not Localize Around the Phagosome Membrane in the Absence of RAB-35. Related to Figure 3. (A-C) Localization of YFP-RAB-35 (top) and CTNS-1-mKate2 (middle) within the U.l/rp cells examined during linker cell (LC) death and degradation in the wild type. Bottom: merge, YFP-RAB-35, green; CTNS-mKate2, magenta. Scale bar, 5 μm. Caret, minimal colocalization between RAB-35 and CTNS-1. (D-F) Imaging details as in (A), except that mCh-RAB-5 is imaged in a *rab-35*(*b1013*) animal (top). Bottom: mCh-RAB-5 overlaid on DIC image. Arrowhead, linker cell.

**Figure S6.**
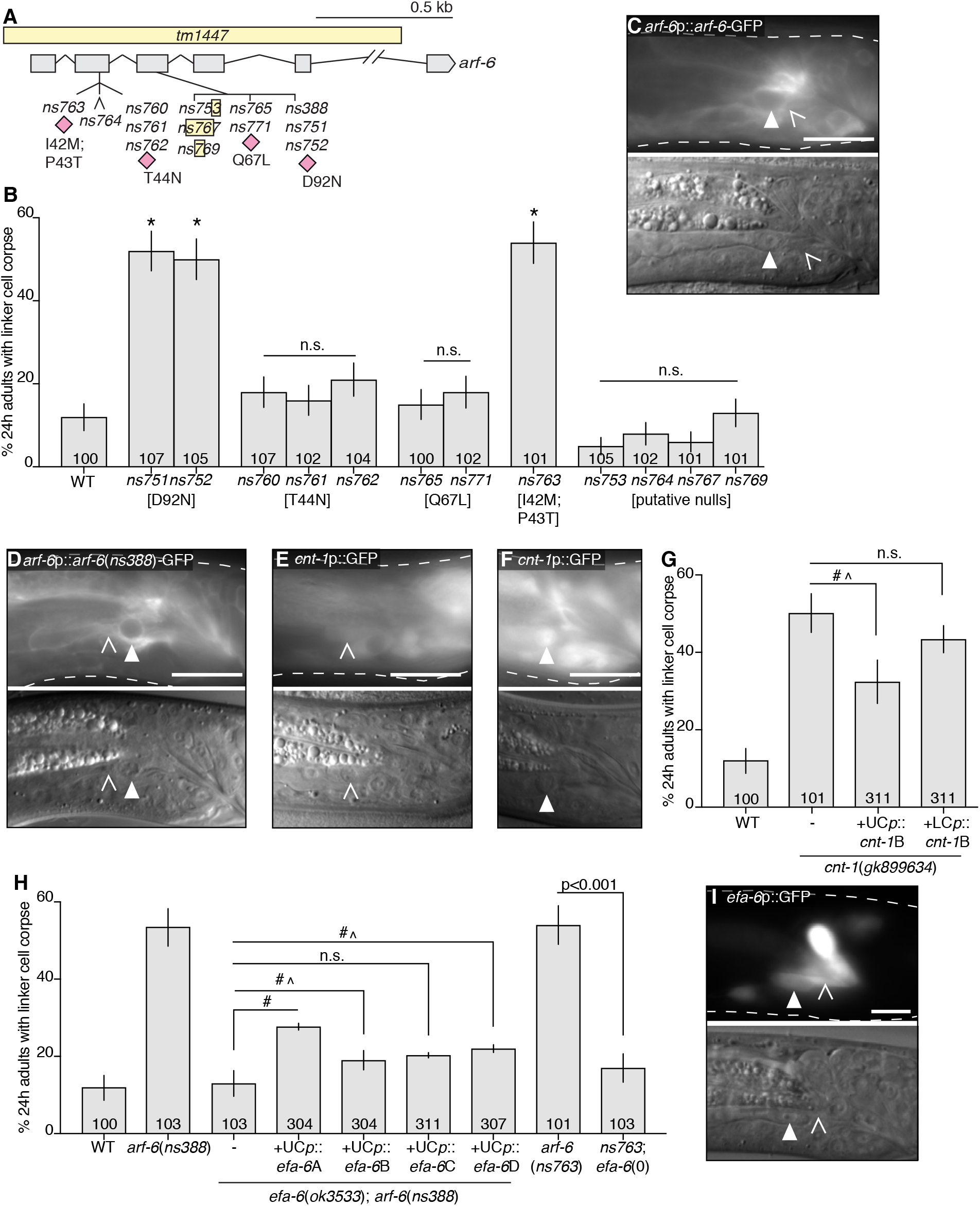
CNT-1/ACAP2 and EFA-6/EFA6 Regulate ARF-6, Related to Figure 4 and Figure 5. (A) *arf-6* genomic locus. Yellow box, deletion alleles. Caret, single nucleotide insertion allele *ns764*. Pink diamonds, point mutations. (B) Histogram details as in Figure 2B. (C) *arf-6* translational reporter expression. Arrowhead, linker cell. Caret, U.lp or U.rp cell. Top: GFP; bottom: DIC. Scale bar, 10 μm. (D-F) Expression reporter for *arf-6*(*ns388*), and *cnt-1*. (G) Histogram as in B. UCp = *lin-48*p. LCp = *mig-24*p. Error bars represent standard error of the proportion or standard error of the mean. #, significantly different from both *cnt-1*(*gk899634*) and wild type.^, only two lines out of three show significant rescue. (H) Histogram as in B. UCp = *lin-48*p. LCp = *mig-24*p. Error bars represent standard error of the proportion or standard error of the mean. #, significantly different from both *efa-6*(*ok3533*); *arf-6*(*ns388*) and *arf-6*(*ns388*).^, only one line out of three show significant rescue. (I) *efa-6* transcriptional reporter. Image information as in C.

**Figure S7.**
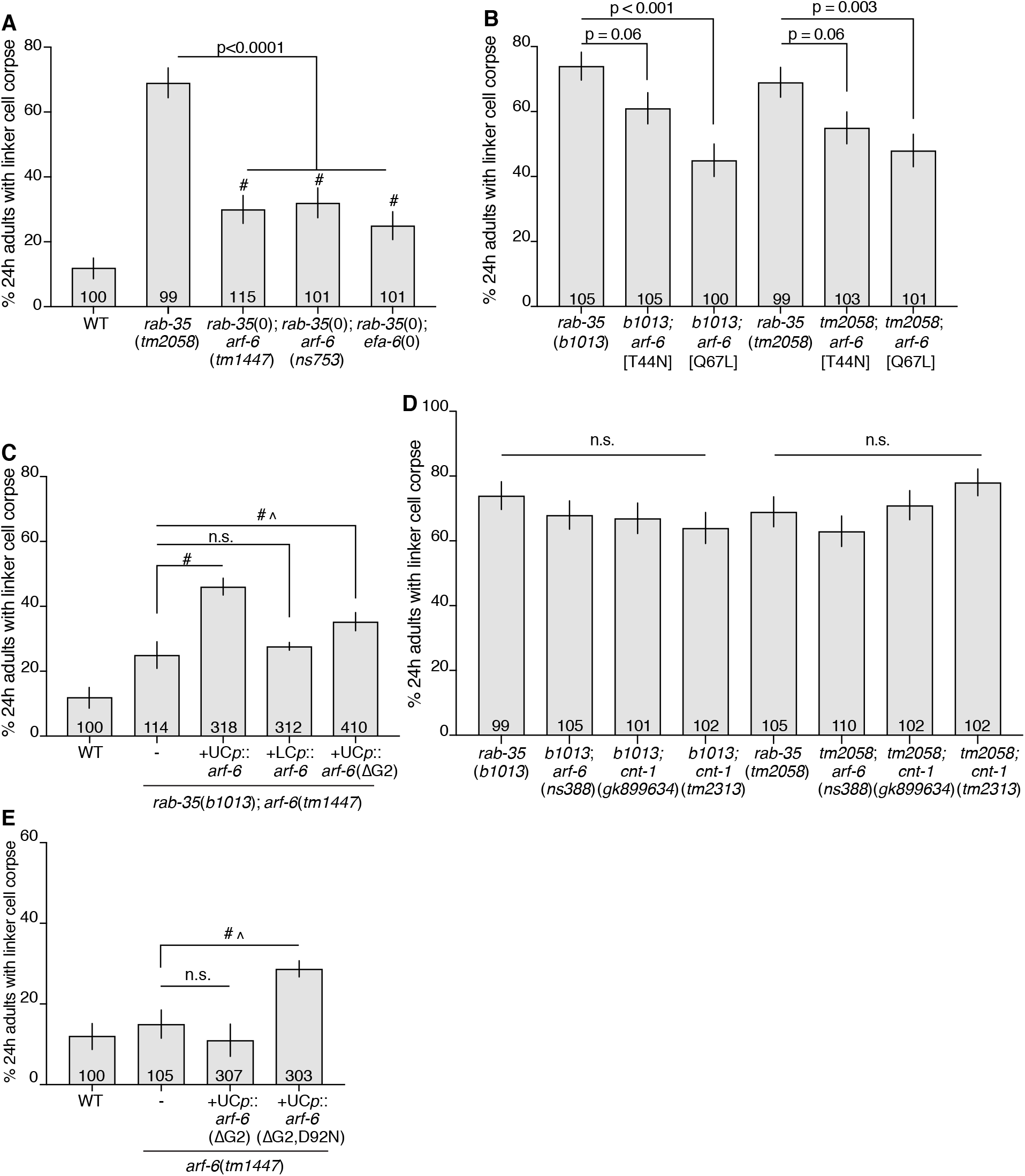
RAB-35 and ARF-6 Are in the Same Genetic Pathway, Related to Figure 6. (A-E) Histogram details as in Figure 2B. ΔG2 = deletion of glycine2, necessary for myristoylation. #, (A) significantly different from single mutants, (C) significantly different from both *rab-35*(*b1013*); *arf-6*(*tm1447*) and *rab-35*(*b1013*), or (E) significantly different from both *arf-6*(*tm1447*) and *arf-6*(*ns388*).^, only 2/4 lines showed significant rescue.

## Supplementary Movie Legends

**Movie S1. Wild type linker cell death and degradation, Related to Figure 1**. Strain contains a linker cell reporter pseudocolored green (*mig-24*p::Venus), a U.l/rp cell reporter pseudocolored magenta (*lin-48*p::mKate2), and *him-5*(*e1490*). Time, hours:minutes after first contact. Same animal as Figure 1.

**Movie S2. *rab-35*(*b1013*) linker cell degradation is delayed, Related to Figure 2**. Strain contains a linker cell reporter pseudocolored green (*mig-24*p::Venus), a U.l/rp cell reporter pseudocolored magenta (*lin-48*p::mKate2), and *him-5*(*e1490*). Time, hours:minutes after first contact. Same animal as in Figure 2G, K.

**Movie S3. *arf-6*(*ns388*) linker cell degradation is delayed, Related to Figure 4**. Strain contains a linker cell reporter pseudocolored green (*mig-24*p::Venus), a U.l/rp cell reporter pseudocolored magenta (*lin-48*p::mKate2), and *him-5*(*e1490*). Time, hours:minutes after first contact. Animal same as in Figure 4C.

**Table S1.**
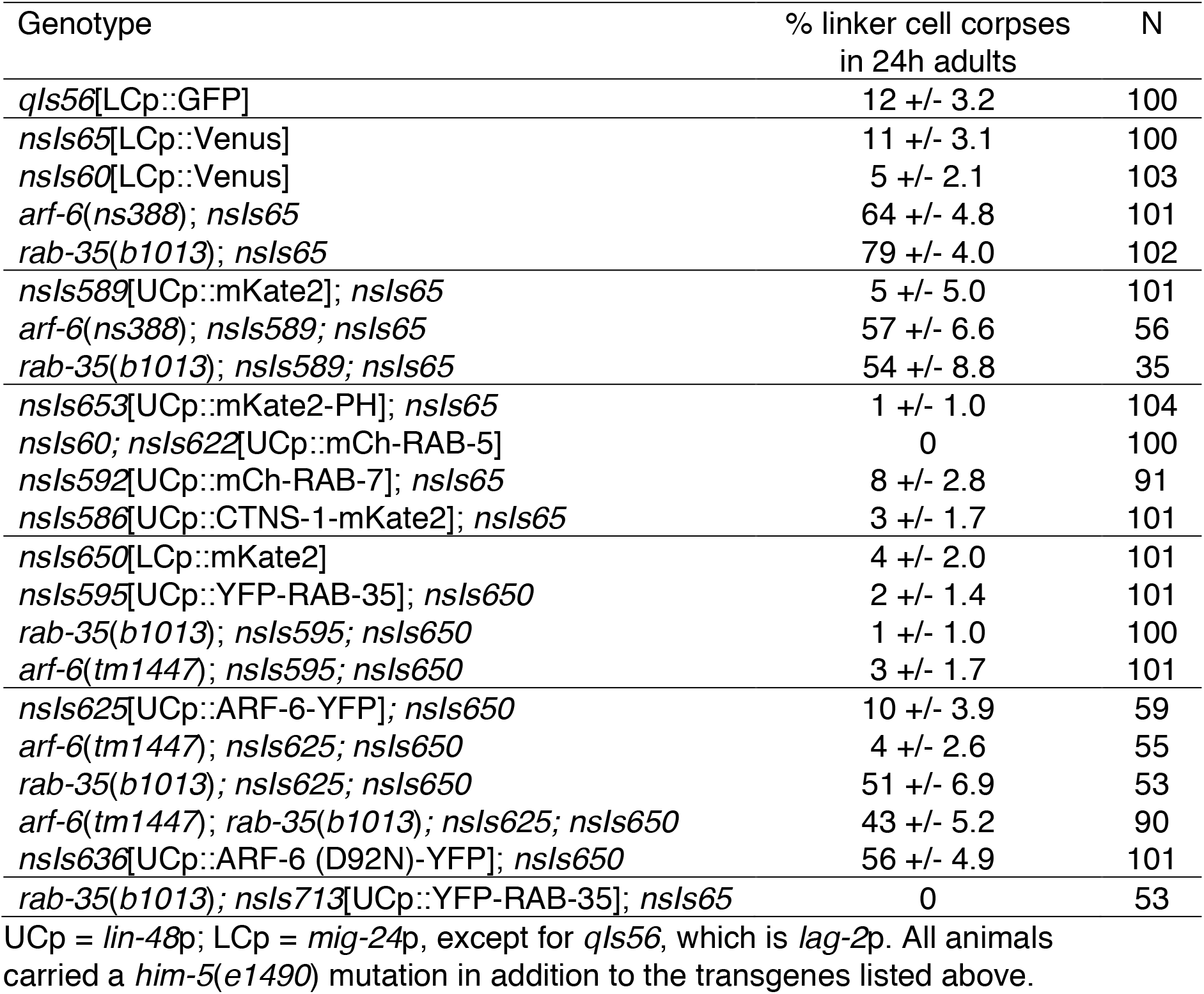
Reporter identity does not affect linker cell degradation (mean +/− SE of proportion). Related to Figures 1–4,6,S5.

| Genotype | % linker cell corpses<br>in 24h adults | N |
| --- | --- | --- |
| *qls56*[LCp::GFP] | 12 +/- 3.2 | 100 |
| *nsIs65*[LCp::Venus] | 11 +/- 3.1 | 100 |
| *nsIs60*[LCp::Venus] | 5 +/- 2.1 | 103 |
| *arf-6(ns388); nsIs65* | 64 +/- 4.8 | 101 |
| *rab-35(b1013); nsIs65* | 79 +/- 4.0 | 102 |
| *nsIs589*[UCp::mKate2]; *nsIs65* | 5 +/- 5.0 | 101 |
| *arf-6(ns388); nsIs589; nsIs65* | 57 +/- 6.6 | 56 |
| *rab-35(b1013); nsIs589; nsIs65* | 54 +/- 8.8 | 35 |
| *nsIs653*[UCp::mKate2-PH]; *nsIs65* | 1 +/- 1.0 | 104 |
| *nsIs60; nsIs622*[UCp::mCh-RAB-5] | 0 | 100 |
| *nsIs592*[UCp::mCh-RAB-7]; *nsIs65* | 8 +/- 2.8 | 91 |
| *nsIs586*[UCp::CTNS-1-mKate2]; *nsIs65* | 3 +/- 1.7 | 101 |
| *nsIs650*[LCp::mKate2] | 4 +/- 2.0 | 101 |
| *nsIs595*[UCp::YFP-RAB-35]; *nsIs650* | 2 +/- 1.4 | 101 |
| *rab-35(b1013); nsIs595; nsIs650* | 1 +/- 1.0 | 100 |
| *arf-6(tm1447); nsIs595; nsIs650* | 3 +/- 1.7 | 101 |
| *nsIs625*[UCp::ARF-6-YFP]; *nsIs650* | 10 +/- 3.9 | 59 |
| *arf-6(tm1447); nsIs625; nsIs650* | 4 +/- 2.6 | 55 |
| *rab-35(b1013); nsIs625; nsIs650* | 51 +/- 6.9 | 53 |
| *arf-6(tm1447); rab-35(b1013); nsIs625; nsIs650* | 43 +/- 5.2 | 90 |
| *nsIs636*[UCp::ARF-6 (D92N)-YFP]; *nsIs650* | 56 +/- 4.9 | 101 |
| *rab-35(b1013); nsIs713*[UCp::YFP-RAB-35]; *nsIs65* | 0 | 53 |

UCp = *lin-48p*; LCp = *mig-24p*, except for *qls56*, which is *lag-2p*. All animals carried a *him-5(e1490)* mutation in addition to the transgenes listed above.

**Table S2.**
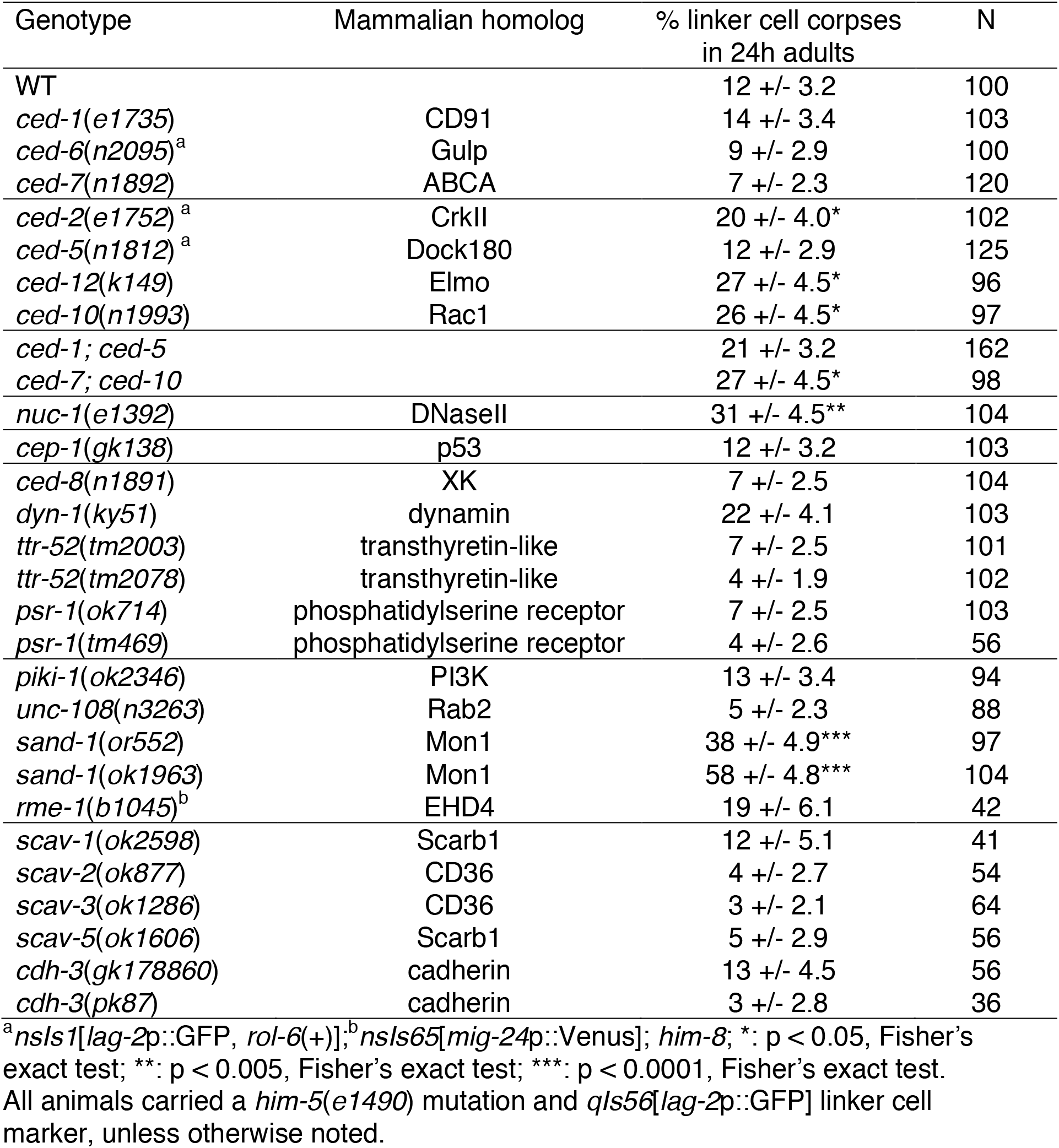
Canonical apoptotic engulfment genes play a minor role in linker cell corpse removal (mean +/− SE of proportion). Related to Figure 1.

**Table S3.** Other putative ARF-6 GAPs have no linker cell defect. Related to Figure 4.

| Genotype | % linker cell corpses<br>in 24h adults | N | homolog |
| --- | --- | --- | --- |
| WT | 12 +/- 3.2 | 100 |  |
| *F23H11.4(gk585084)* | 8 +/- 4.3 | 39 | ARAP1-3/ ArfGAP |
| *F23H11.4(gk165637)* | 13 +/- 3.4 | 96 | ARAP1-3/ ArfGAP |
| *git-1(ok1848)* | 10 +/- 3.9 | 59 | GIT1 |
| *git-1(gk392605)* | 15 +/- 5.6 | 41 | GIT1 |
| *gap-2(ok1001)* | 10 +/- 4.2 | 52 | SynGAP/RasGAP |
| *cnt-2(gm377)* | 12 +/- 3.2 | 102 | ArfGAP |

All animals carried a *him-5(e1490)* mutation and *qls56[lag-2p::GFP]* linker cell marker.

**Table S4.** Mutations in the canonical apoptotic engulfment genes *ced-1* and *ced-5* do not enhance mutations in *rab-35*.

| Genotype | % linker cell corpses in 24h<br>adults | N |
| --- | --- | --- |
| WT | 12 +/- 3.2 | 100 |
| *ced-1(e1735)* | 14 +/- 3.4 | 103 |
| *ced-5(n1812)<sup>a</sup>* | 12 +/- 2.9 | 125 |
| *rab-35(b1013)* | 74 +/- 4.2 | 105 |
| *rab-35(tm2058)* | 69 +/- 4.6 | 99 |
| *ced-1; rab-35(b1013)* | 77 +/- 4.2 | 101 |
| *ced-5; rab-35(b1013)* | 63 +/- 4.8 | 101 |
| *ced-1; ced-5; rab-35(b1013)* | 62 +/- 4.8 | 103 |
| *ced-1; rab-35(tm2058)* | 56 +/- 4.9 | 102 |
| *ced-5; rab-35(tm2058)* | 58 +/- 4.9 | 102 |
| *ced-1; ced-5; rab-35(tm2058)* | 63 +/- 4.7 | 104 |

<sup>a</sup> = *nsIs1*[*lag-2p::GFP*, *rol-6(+)*]. All animals carried a *him-5(e1490)* mutation and *qls56*[*lag-2p::GFP*] linker cell marker, unless otherwise noted.

## Materials and Methods

### Contact for Reagent and Resource Sharing

Further information and requests for reagents may be directed to, and will be fulfilled by the corresponding author Shai Shaham.

### Experimental Model and Subject Details

#### C. elegans

*C. elegans* strains were raised at 20°C on nematode growth medium (NGM) seeded with OP50 bacteria, unless otherwise indicated. Wild-type animals were of the Bristol N2 strain. All strains have one of two mutations that generate a high incidence of male progeny, *him-8*(*e1489*) IV or *him-5*(*e1490*) V. Mutants recovered by EMS mutagenesis were outcrossed five times before use. Transgenic lines were generated by injection of plasmid DNA mixes into the hermaphrodite gonad. Integrated transgenic strains were generated with UV/ trioxalen treatment^55^ (Sigma, T6137) and were outcrossed at least four times before imaging experiments. Most strains also had one of three integrated linker cell markers qIs56[*lag-2*p::GFP] V, nsIs65[*mig-24*p::Venus] X, or nsIs650[*mig-24*p::mKate2] X. Other alleles and transgenes are as follows:

LGI: *rab-2*(*n3263*), *ced-1*(*e1735*), *cep-1*(*gk138*), *dpy-5*(*e907*)
LGII: *cnt-1*(*gk899634*), *cnt-1*(*tm2313*)
LGIII: *rab-35*(*b1013*), *rab-35*(*tm2058*), *tbc-10*(*gk388086*), *unc-119*(*ed3*), *ttr-52*(*tm2003*), *ttr-52*(*tm2078*), *F23H11.4*(*gk585084*), *F23H11.4*(*gk165637*), *cnt-2*(*gm377*)
LGIV: *arf-6*(*tm1447*), *arf-6*(*ns388*), *arf-6*(*ns751*[*D92N*]), *arf-6*(*ns752*[D92N]), *arf-6*(*ns753*), *arf-6*(*ns760*[T44N]), *arf-6*(*ns761*[T44N]), *arf-6*(*ns762*[T44N]), *arf-6*(*ns763*[I42M;P43T]), *arf-6*(*ns765*[Q67L]), *arf-6*(*ns771*[Q67L]), *arf-6*(*ns764*), *arf-6*(*ns767*), *arf-6*(*ns769*), *sand-1*(*or552*), *sand-1*(*ok1963*), *efa-6*(*ok3533*), *ced-10*(*n1993*), *ced-5*(*n1812*), *psr-1*(*ok714*), *psr-1*(*tm469*)
LGX: *rme-4*(*ns410*), *rme-4*(*ns412*), *rme-4*(*b1001*), *rme-4*(*tm1865*), *tbc-13*(*ok1812*), *piki-1*(*ok2346*), *him-4*(*e1267*), *git-1*(*ok1848*), *git-1*(*gk392605*), *gap-2*(*ok1001*)

#### Yeast two-hybrid assays

The yeast strain NMY51 (Dualsystems Biotech) was maintained at 30°C on YPAD plates. Yeast were plated on SD-Leu-Trp- after transformation, and 5μl of OD600 = 0.4 of transformants were spotted on SD-Leu-Trp- His- to test for interactions.

### Method Details

#### Forward genetic screen and artificial insemination

Young L4 hermaphrodites containing qIs56 [*lag-2*p::GFP] and *him-5*(*e1490*) were mutagenized with 75mM ethylmethanesulfonate (EMS, Sigma M0880) for 4h at 20°C. Animals were synchronized twice by bleaching, and male animals were enriched by passing through a 40μm cell strainer (BD Falcon) passively for 20 min^56^. Male animals fit more readily through the mesh than hermaphrodites and were enriched to ~88%. Animals were then scored under a fluorescent dissecting microscope (Leica) for presence of a linker cell by GFP. GFP-expressing animals were isolated away from hermaphrodites for 24h to accumulate spermatids within the gonad. Mutant males were immobilized on a dried agar pad under immersion oil, and spermatids removed with a needle containing SM buffer (50 mM HEPES, pH 7, 50 mM NaCl, 25 mM KCl, 5 mM CaCl2, 1 mM MgSO4, 1 mg/ml BSA)^28^. An N2 hermaphrodite with a single row of eggs was then immobilized on the same dried agar pad and gently injected with the collected sperm through the vulva. We typically tried to insert all spermatids into a single hermaphrodite. GFP+ heterozygous progeny were collected, allowed to self fertilize, and linker cell persistence confirmed under a dissecting microscope.

#### Gene identification

A combination of Hawaiian Snip-SNP mapping^57^ and whole genome sequencing was using to identify *rme-4*(*ns410*), *rme-4*(*ns412*), and *arf-6*(*ns388*)^58^ using output from galign^59^. *rme-4* was confirmed by fosmid rescue using WRM0615bE09, and *arf-6* identification was confirmed by mutant cDNA expression and CRISPR alleles.

#### Linker cell survival and corpse persistence assays

Linker cell death was scored as previously described^21^. Briefly, unstarved gravid hermaphrodites were bleached to isolate embryos, which were allowed to hatch overnight in M9. Synchronized L1 animals were released on 9-cm NGM plates seeded with OP50 or HT115 *E. coli* containing the RNAi construct of interest on IPTG-RNAi plates. Male animals were isolated onto a new plate prior to the L4-to-adult transition based on full retraction of the male tail tip with rays visible under the unshed L4 cuticle. 24h later, these animals were mounted onto 2% agarose-water pads, anaesthetized in 25 mM sodium azide, and examined on an Axioplan 2 fluorescence microscope (Zeiss) with a 63x/1.4NA objective (Zeiss) and Nomarski optics. The linker cell was identified by location, morphology, and fluorescence from reporter transgenes. Cells were then scored as surviving, dead, or gone based on DIC morphology and fluorescence.

#### Generation of *arf-6* alleles using CRISPR-Cas9 genome editing

Alleles of *arf-6* were generated using the co-CRISPR-based genome editing method as previously described^60^. pDD162 was used as a vector backbone, and the following sgRNA sequences were added for each individual CRISPR attempt (D92N: 5’ CTAAAAACCTGTCTCTATCAG 3’; T44N: 5’ TCAGTGACCACAATGACGA 3’; Q67L 5’ CTTCAGGACGTCGGCGGAC 3’) to generate *arf-6* targeting vectors. A *dpy-10* sgRNA-pDD162-based vector was also generated (5’ GCTACCATAGGCACCACGAG 3’). Single-stranded repair oligos were PAGE purified and ordered from Sigma (D92N: 5’ aggaaaaaaaccaatttttccgcatttttcgcctaaaaacCTGTCTCTATtAGCaGCGTCCATCACAAAAATGAGCGCCTGAGTTCCTGTGTAATAATGTC 3’; T44N: 5’ agCAATTCTGTACAAACTGAAGCTCGGGCAATCAGTGACCACAATTCCGAacGTGGGCTTCAATGTGGAGACTGTCACGTATAAAAATATCAAATTCAACGT 3’; Q67L 5’ ggcctaaaaacccccaaaaaccccaatttttcttcagGACGTCGGCGGACttGACAAAATTCGACCCCTCTGGCGACATTATTACACAGGAACTCAGGCGCT 3’, dpy-10(cn64): 5’ CACTTGAACTTCAATACGGCAAGATGAGAATGACTGGAAACCGTACCGCATGCGGTGCCTATGGTAGCGGAGCTTCACATGGCTTCAGACCAACAGCCTAT 3’^60^. N2 animals were injected with the following mix: 50 ng/μl *dpy-10* sgRNA, 50 ng/μl *arf-6* targeting vector, 20 ng/μl *dpy-10*(*cn64*) repair oligo, 20 ng/μl *arf-6* repair oligo in 1x injection buffer (20mM potassium phosphate, 3mM potassium citrate, 2% PEG, pH 7.5). F1 animals with Dpy or Rol phenotypes were picked to individual plates, indicating a CRISPR-based editing event had occurred. F1 animals were allowed to lay eggs, and then genotyped for successful co-conversion of the *arf-6* locus using PCR and restriction enzyme screening or Cel1 digestion of heteroduplex DNA^61^. Non-Rol, non-Dpy F2 animals were then singled and homozygosed for the *arf-6* mutation.

#### Germline transformation and rescue experiments

Plasmid mixes containing the plasmid of interest, co-injection markers, and pBluescript were injected into both gonads of young adult hermaphrodites^62^. Injected animals were singled onto NGM plates and allowed to grow for two generations. Transformed animals based on co-injection markers were picked onto single plates, and screened for stable inheritance of the extrachromosomal array. Only lines from different P0 injected hermaphrodites were considered independent. For cell-specific rescue experiments, animals expressing mCherry in either the linker cell or the U.l/rp cells were picked to a new plate at the early L4 stage, before linker cell death, to avoid bias. Isolated animals were then staged based on tail morphology under a white light microscope for appropriate linker cell scoring. Transgenes used are listed in the following table.

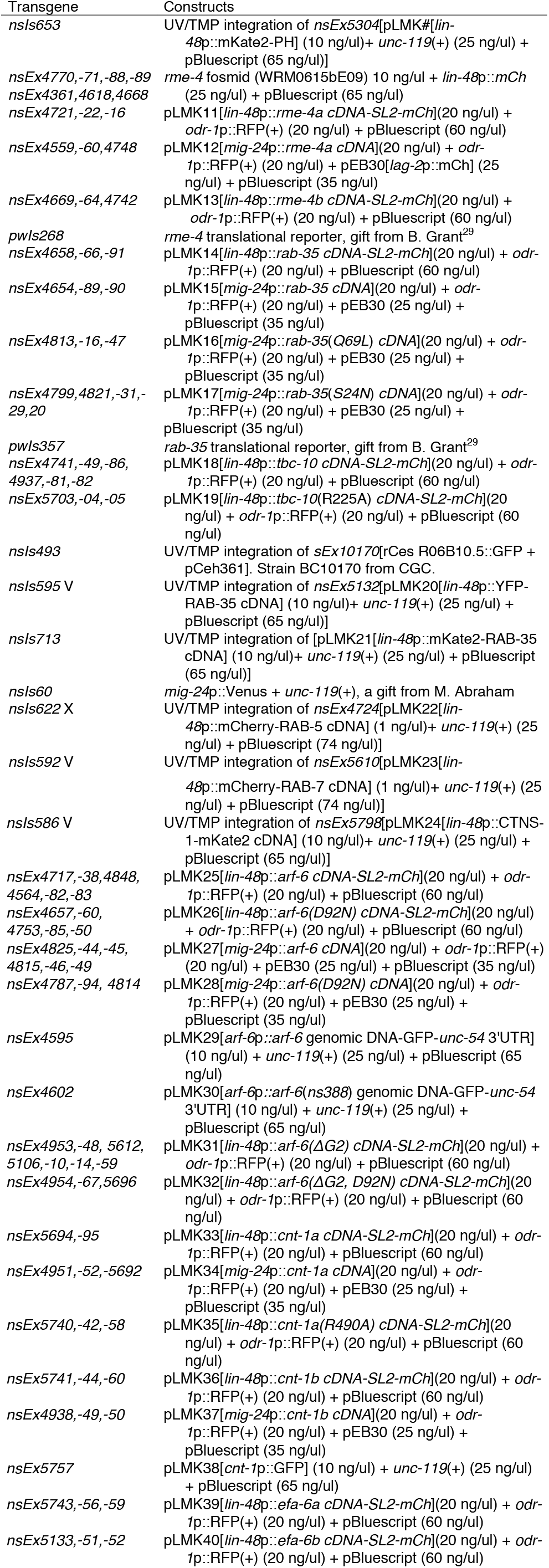
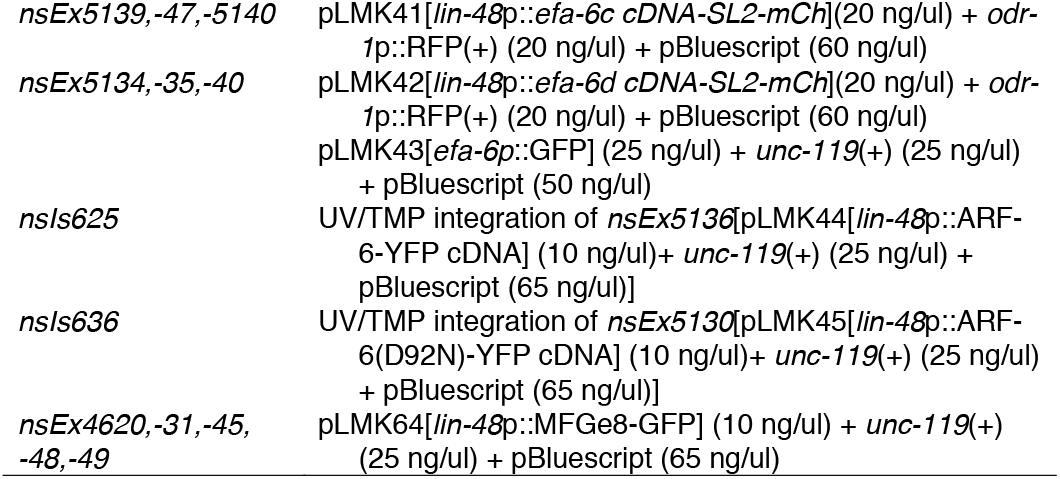

#### Plasmid construction

Plasmids containing *lin-48p*, *mig-24*p, or transcriptional reporters were cloned using multi-piece one-step cloning into a modified pPD95.75 backbone lacking GFP. For CRISPR-related vectors, plasmids were generated using a site-directed plasmid mutagenesis protocol on pDD162^63,64^. Yeast vectors were generated using traditional cloning from either a pLexA-N-terminus vector or a pGAL4AD-N-terminus vector (Dualsystems Biotech), and single amino acid change variants were generated using QuikChange (Agilent).

Additional Plasmids
| Plasmid | Details |
| --- | --- |
| pLMK46 | LexA-CNT-1A, Yeast two-hybrid (Y2H) |
| pLMK47 | LexA-CNT-1B, Y2H |
| pLMK48 | GAD-ARF-6, Y2H |
| pLMK49 | GAD-ARF-6(D92N), Y2H |
| pLMK50 | GAD-ARF-6(I42M,P43T), Y2H |
| pLMK51 | LexA-ARF-6(T44N), Y2H |
| pLMK52 | GAD-EFA-6A, Y2H |
| pLMK53 | GAD-EFA-6B, Y2H |
| pLMK54 | GAD-EFA-6C, Y2H |
| pLMK55 | GAD-EFA-6D, Y2H |
| pLMK56 | LexA-RAB-35(S24N), [GDP], Y2H |
| pLMK57 | LexA-RAB-35(Q69L), [GTP], Y2H |
| pLMK58 | GAD-CNT-1A, Y2H |
| pLMK59 | GAD-CNT-1B, Y2H |
| pLMK60 | GAD-RME-4A, Y2H |
| pLMK61 | GAD-RME-4B, Y2H |
| pLMK62 | *arf-6*(D92N) CRISPR targeting vector, pDD162 backbone |
| pLMK63 | *arf-6*(T44N) CRISPR targeting vector, pDD162 backbone |
| pLMK64 | *arf-6*(Q67L) CRISPR targeting vector, pDD162 backbone |

#### Protein structures

The co-crystal structure of human Arf6 with the ArfGAP ASAP3 (PDB 3LVQ) was examined in Pymol (The PyMOL Molecular Graphics System, Version 1.2r3pre, Schrödinger, LLC.) in silico mutagenesis for D92N and I42M;P43T was performed and hydrophilic interactions tested within Pymol. Key residues noted in Figure 5A-D in ASAP3 are conserved in *C. elegans* CNT-1, and those for Arf6 are conserved in *C. elegans* ARF-6.

#### Long-term imaging and movie generation

Early L4 male animals were imaged in a microfluidic device described in detail in^24^. Animals were fed a constant flow of NA22 bacteria in S medium, supplemented with kanamycin (50ng/μl) to prevent bacterial overgrowth. Animals were immobilized and imaged every 8 minutes for at least 20h. Mutant animals were typically imaged for >30h. Exposure time and light intensity was held constant across strains when the same integrated transgene was imaged. Occasionally tail development was perturbed by the flow of medium and repeated immobilization procedures, and these animals were not included in subsequent analyses.

#### Deconvolution

Images were cropped to a region of interest surrounding the linker cell. We measured the point-spread function (PSF) of our optical setup using red (580/605nm) and green (505/515nm) fluorescent 200nm beads (ThermoFisher Scientific). Using these PSFs, we deconvolved the cropped fluorescence z-stacks using the classic maximum likelihood estimation (CMLE) algorithm with standard parameters (refractive index of imaging medium: 1.338) in the Huygens Essential software (Huygens Essential 3.7.1) by Scientific Volume Imaging (SVI).

#### Image Analysis

Image analysis was performed using custom-written Matlab R2016b (Mathworks) scripts. To overlay imaging frames, we straightened each three-dimensional image stack using a previously published algorithm^24,65^ based on a manually selected worm backbone in the DIC channel. To correct for small residual animal movements during multi-channel acquisition, obvious landmarks visible in the fluorescence channels, such as the cloaca, fluorescently labeled cells in the animal’s tail, or vesicles within the U-cell descendants were then manually aligned to the DIC channel in each frame. Time-lapse movies were generated by centering all straightened, aligned images on the linker cell and cropping the entire movie to an appropriate size. Frames were removed from the movies whenever residual animal movement excessively blurred the images.

#### RNAi assay

pL4440 RNAi plasmids containing coding sequence to *rme-4* or *tbc-7* were isolated from the Ahringer Library^66^ (Source BioScience) and the coding sequence was confirmed by sequencing. These plasmids were grown at 37°C in HT115 bacteria, which have a plasmid containing an inducible promoter driving T7 RNA polymerase. The bacteria were plated on NGM plates that had been supplemented with the antibiotic carbomycin and IPTG. One day later, synchronized L1 animals were plated on the RNAi plates and grown for two days before assaying for linker cell corpse persistence.

#### Yeast two-hybrid assay

Strain and vectors used in assay were from the DUALHybrid Kit (Dualsystem Biotech). Bait cDNA was cloned into the pLexA plasmid and prey cDNA was cloned into pGAD vectors. These plasmids were cotransformed into NMY51 yeast strain using the lithium acetate method described in the DualHybrid manual. Selection was performed on SD plates lacking the amino acids leucine, tryptophan, and histidine.

#### Apoptotic corpse assay

We counted the number of apoptotic corpses in 3-fold embryos mounted on agar pads and examined at 63x using DIC (Nomarski optics). These corpses could be distinguished by their high refractility.

#### Quantification and Statistical Analysis

Statistical analysis was performed using GraphPad Prism. Statistical parameters including mean ± standard error of the proportion, mean ± SEM, and N are reported in the main text, figures and figure legends. Data is judged to be statistically significant when p < 0.05 by Fisher’s exact test, χ2 test, or student’s T test, where appropriate.

#### Data Analysis

When comparing a transgenic line to the parental strain, a minimum of three independent lines was scored. Approximately 100 animals from each of these lines were examined for linker cell defects, and compared to the parental strain using a Fisher’s exact test to determine significance. For ease of presentation, we pooled the lines from a given experiment and displayed them on the relevant figure using mean ± SEM (N = 3, 4, or 5 lines). If only a subset of the three lines showed significance using a Fisher’s exact test, we indicated this on the figure and figure legend.

